# Inference and validation of an integrated regulatory network of autism

**DOI:** 10.1101/2020.06.08.139733

**Authors:** Maedeh Ganji, Modjtaba Emadi-Baygi, Maryam Malek, Parvaneh Nikpour

## Abstract

Autism is a complex neurodevelopmental disorder. Functional roles of several non-coding transcripts including long noncoding RNAs (lncRNAs) have been shown to influence the pathobiology of autism. We hypothesized that there are more autism-associated lncRNAs to be discovered. Here, we utilized a systems biology approach to identify novel lncRNAs that might play a role in the molecular pathogenesis of autism. Based on the data provided by the Simons Foundation Autism Research Initiative (SFARI), a three-component regulatory network comprising mRNAs, microRNAs (miRNAs) and lncRNAs was constructed. Functional enrichment analysis was performed to identify molecular pathways potentially mediated by components of the network. The potential association of four candidate lncRNAs with autism was investigated experimentally by developing and verifying a valproic acid (VPA)-exposed mouse model of autism. We composed a network of 33 mRNA, 25 miRNA and 4 lncRNA nodes associated with neurologically-relevant pathways and functions. We then verified the differential expression of four candidate lncRNAs: *Gm10033*, *1500011B03Rik*, *A930005H10Rik* and *Gas5* in the brain of VPA-exposed mice. We furthermore identified a novel splice variant of *Gm10033*, designated as *Gm10033-*Δ*Ex2*, which was expressed in various mouse tissues. The integrative approach, we utilized, combines the analysis of a three-component regulatory network with experimental validation of targets in an animal model of autism. As a result of the analysis, we prioritized a set of candidate autism-associated lncRNAs. These links add to the common understanding of the molecular and cellular mechanisms that are involved in disease etiology, specifically in the autism.

## 1. Introduction

Autism is a heterogeneous, heritable neurodevelopmental disorder with an onset in early childhood (1, 2) and is characterized by deficits in the fields of social communication, social interaction as well as repetitive behaviors (3). According to the reports of The Centers for Disease Control and Prevention (CDC), autism prevalence rates have increased from 1 in 200 in the late 1990s to 1 in 59 children aged 8 years in 2014 (4, 5).

Due to the complex nature of autism, it is challenging to unravel its molecular disease mechanisms at different levels of functional organization. Recent studies have identified hundreds of autism-related mRNAs and microRNAs (miRNAs) by applying genome-scale expression analysis (6–8). Furthermore, a novel class of regulatory RNAs, long non-coding RNAs (lncRNAs) was shown to implicate in several neurodevelopmental disorders, including autism (9). lncRNAs are a group of non-coding RNAs longer than 200 nucleotides. They are involved in a range of cellular/molecular processes, namely, transcription, translation, and post-transcriptional RNA regulation, and they play pivotal roles in neurobiological and neuropathological conditions (10).

Autism likely involves dysregulation of multiple coding and non-coding transcripts related to brain function and development. Several gene expression profiling studies have demonstrated aberrant expression of coding (11) and non-coding RNAs including miRNAs (6) and lncRNAs (12) in autistic brains. These data suggest that non-coding regions of the genome may have a major impact on the autism molecular pathology (12).

Given the accumulated data on autism, it is now possible to implement bioinformatics and systems biology pipelines, which can predict potential disease-gene associations and eventually lead to a better understanding of the molecular aspects of autism. Thus far, several studies have investigated gene expression regulatory networks comprising of mRNAs, miRNAs, or lncRNAs in the peripheral blood or brain tissue of autistic patients. For example, Huang et al (7) constructed an integrated miRNA regulatory network presenting the relationships between the miRNAs, mRNAs, and miRNA transcription factors. To integrate the genome-wide-expressed lncRNAs with the synaptic mRNAs in autism spectrum disorder, Wang et al (13) constructed and analyzed such a network to associate differential gene expression with gene ontology, biological pathways, and the regulatory functions of the lncRNAs. While previous studies have constructed two-component regulatory networks of mRNAs, miRNAs, and lncRNAs to identify autism-associated genes, none of these studies have concurrently explored the relationships among these three diverse RNA species in the context of an integrated multilayered regulatory network.

Therefore, the purpose of this study was to find novel lncRNAs that might play a role in the molecular pathogenesis of autism. To do so, we retrieved autism-related mRNAs from the Simons Foundation Autism Research Initiative (SFARI), the miRNAs which target them, as well as related lncRNAs. Then, we mapped them onto a three-component regulatory network. Moreover, we developed a valproic acid (VPA)-exposed mouse model of autism to experimentally validate the association of selected lncRNAs with autism. We furthermore describe the identification and characterization of a novel alternatively spliced *Gm10033* transcript lacking exon 2 and designated it as *Gm10033-*Δ*Ex2.* The sequence was submitted to the GenBank database under the accession number: MN231628.1.

## 2. Materials and Methods

### 2.1. Data collection and network construction

A list of 367 autism susceptibility genes was obtained from the Gene’s Animal Models Module of SFARI (14) on 19 September 2018. The list was then filtered to contain only the 113 *Mus musculus* genes with SFARI gene rankings 1–4 (high, strong candidate, suggestive and minimal evidence) and genes with more than one animal model available. Of note, in the latest update of SFARI on 2020 Q3, all genes are organized into 3 categories (high, strong candidate, and suggestive evidence) that confirms the high association of these genes with autism. An interaction network was then constructed and visualized using the CluePedia plug-in (version 1.3.5) in Cytoscape (version 3.2.1) (15, 16). The STRING Action File, which is available from CluePedia, was the source for action types including activation, inhibition, binding, and post-translational modification. Then, miRNAs targeting the top 20% selected genes, sorted based on the degree centrality, were retrieved from miRTarBase (17) and TargetScan (18). We built a two-component regulatory network including mRNAs and miRNAs utilizing Cytoscape, and miRanda, mirTarBase, and miRecords of CluePedia plug-in. lncRNAs targeted by the top 20% miRNAs with high degree centrality values were then extracted from LncBase Experimental (Version .2) (19). The result was a three-component regulatory network including mRNAs, miRNAs, and lncRNAs.

### 2.2. Functional enrichment analysis

To identify molecular pathways potentially mediated by mRNAs and miRNAs in the subnetwork (described in the Results’ section), the Kyoto Encyclopedia of Genes and Genomes (KEGG) pathway enrichment of Enrichr (20) and DIANA-mirPath (21) were exploited, respectively. Furthermore, STRING (22) was employed to identify significantly enriched Gene Ontology (GO) terms of subnetwork mRNAs (23).

### 2.3. Protein–protein interaction analysis

A total of 91 mRNAs in the network were analyzed by STRING to predict protein-protein interactions (PPIs). The combined scores greater than 0.9 were selected to construct the PPI network by hiding disconnected nodes. STRING interaction data was further utilized to explore the KEGG pathways retrieved from highly connected proteins.

### 2.4. Phenotype annotations and gene expression profiling of the mouse brain

Phenotype annotations analysis of 91 mRNAs in the network was performed using Disease Ontology (DO) enrichment and Mammalian Phenotype (MP) Ontology Enrichment term annotations from the MGI (Mouse Genome Informatics) resource (24). Any synonyms for each term (gene), which may appear in the literature or other online resources, are added by the database (25). To obtain expression maps of the top 6 genes (*Foxp1*, *Rhno1*, *Tshz3*, *Ctnnb1*, *Grip1*, and *Prkdc*) which were sorted based on the degree centrality in the main network (≥40) of various brain regions within the developing mouse embryo and adult mouse, BrainStars (B*), EMAGE, was exploited.

### 2.5. Animals

Fifty pairs of C57BL/6J mice (Pasteur Institute, Tehran, Iran) were kept in cages with a temperature of 21±1 °C and light conditions (12h:12h, lights at 07:00 a.m.). Mice were given *ad libitum* access to food (standard laboratory pellets) and water. The presence of a vaginal plug after overnight mating indicated successful mating and the date was taken as gestational day 0 (GD 0). At GD 12.5, pregnant females received a single intraperitoneal (i.p.) injection of 500 mg/kg sodium salt of VPA (Sigma, USA) (26), and control pregnant mice were injected with the same volume of physiological saline. All animal experiments were approved by the institutional animal ethics review board of Isfahan University of Medical Sciences. Behavioral tests were conducted between 10 a.m. and 4 p.m. on 8 week-old male pups because of a higher incidence of autism in boys than girls, and more apparent behavioral changes in VPA-exposed males (27, 28).

### 2.6. Behavioral tests

#### 2.6.1. Three-chamber sociability test

Deficits in social communication were assessed with a three-chamber sociability test on male pups. The test was performed in a clear Plexiglass box (20×45×30 cm), with three communicating chambers of equal sizes (29). Two identical wire cages were placed in the left and right chambers. A stranger mouse was placed into one of the wire cages and the subject was allowed to explore freely around the chambers for 10 minutes. Afterward, a novel mouse was placed into the empty wire cage and the subject was allowed to explore the chambers for another 10 minutes (30). The time spent by the subject in each chamber containing the stranger and novel mice was recorded by a video camera (Hangzhou Hikvision Digital Technology Co., Ltd, China). The sociability index (SI) was calculated as the quotient between stranger chamber divided by the empty chamber duration. The social preference index (SPI) was equal to the time spent in the novel divided by the familiar chambers (31).

#### 2.6.2. Open field test

Locomotor activity and anxiety-like behaviors were measured by the open field test (32, 33). The testing chamber was made of white Plexiglass (72×72×50 cm) with a floor divided into 16 squares, while assigning the four central squares as an inner zone. The subject mouse was gently placed in the central area allowing it to explore it undisturbed for 30 minutes. The number of crossings one of the squares with all four paws was an indication of total locomotor activity. Anxiety-like behaviors were assessed by recording the relative time spent in the inner and outer zones. As a sign of repetitive behavior, self-grooming was further recorded and analyzed by a video camera.

#### 2.6.3. Passive avoidance test

Learning and memory defects are the two secondary symptoms of autism that are evaluated by a passive avoidance test. The apparatus comprises two sections: one is a light chamber (25×25×30 cm) and the other one is dark (25×25×30 cm) with a grid floor. For the acquisition trial, the subject mouse was placed into the light chamber. Ten seconds later, the slit door was opened and the latency of the mouse to escape into the dark chamber was recorded. After 10 seconds, a 0.45 mA foot shock was applied for 3 seconds (34). Twenty-four hours later, in the retention trial, the mouse was placed again in the same situation as the previous day, and the latency to escape into the dark chamber was recorded, with no foot shock applied in the retention trial (35).

### 2.7. Tissue collection

At GD 73, male pups were anesthetized via i.p. injection of Ketamine (100 mg/kg; Alfasan Co, Netherlands) and Xylazine (10 mg/kg; Alfasan Co, Netherlands) (36) and decapitated. Brains were immediately removed; the frontal lobe and cerebellum were disparted and rapidly frozen by immersion in the liquid nitrogen and then were stored at −80 °C.

### 2.8. RNA isolation and reverse transcription polymerase chain reaction (RT-PCR)

The tissues were cut into 30-35 mg pieces and homogenized using an automated homogenizer (Precelly^®^24; Bertin Technologies, France) with QIAzol^®^ lysis reagent (Qiagen, USA). The RNA extraction was then performed according to the manufacturer’s instructions. Complementary DNA (cDNA) synthesis was performed on DNase-treated RNA by using random hexamer primers and an M-MLV reverse transcriptase (Yekta Tajhiz Azma, Iran). Quantitative real time RT-PCR was performed using RealQ Plus 2x Master Mix Green High ROX (Ampliqon, Denmark) and specific primers for candidate lncRNAs and hypoxanthine guanine phosphoribosyl transferase (*Hprt*) as a reference gene (37) (**Table 1**) on an Applied Biosystems StepOnePlus™ instrument. The PCR amplification conditions consisted of an initial denaturation at 95°C for 15 minutes, 40 cycles of denaturation at 95°C for 15 seconds, annealing at 60°C for 30 seconds, and extension at 72°C for 30 seconds. Furthermore, to make sure that the actual genes of interest were specifically amplified, the PCR products of some specimens were randomly sequenced with an Applied Biosystems 3500 sequencer (Pishgam Biotech Co., Tehran, Iran). The BioEdit sequence alignment editor (38) and nucleotide BLAST (https://blast.ncbi.nlm.nih.gov/Blast.cgi) were applied to analyze the results.

**Table 1.**
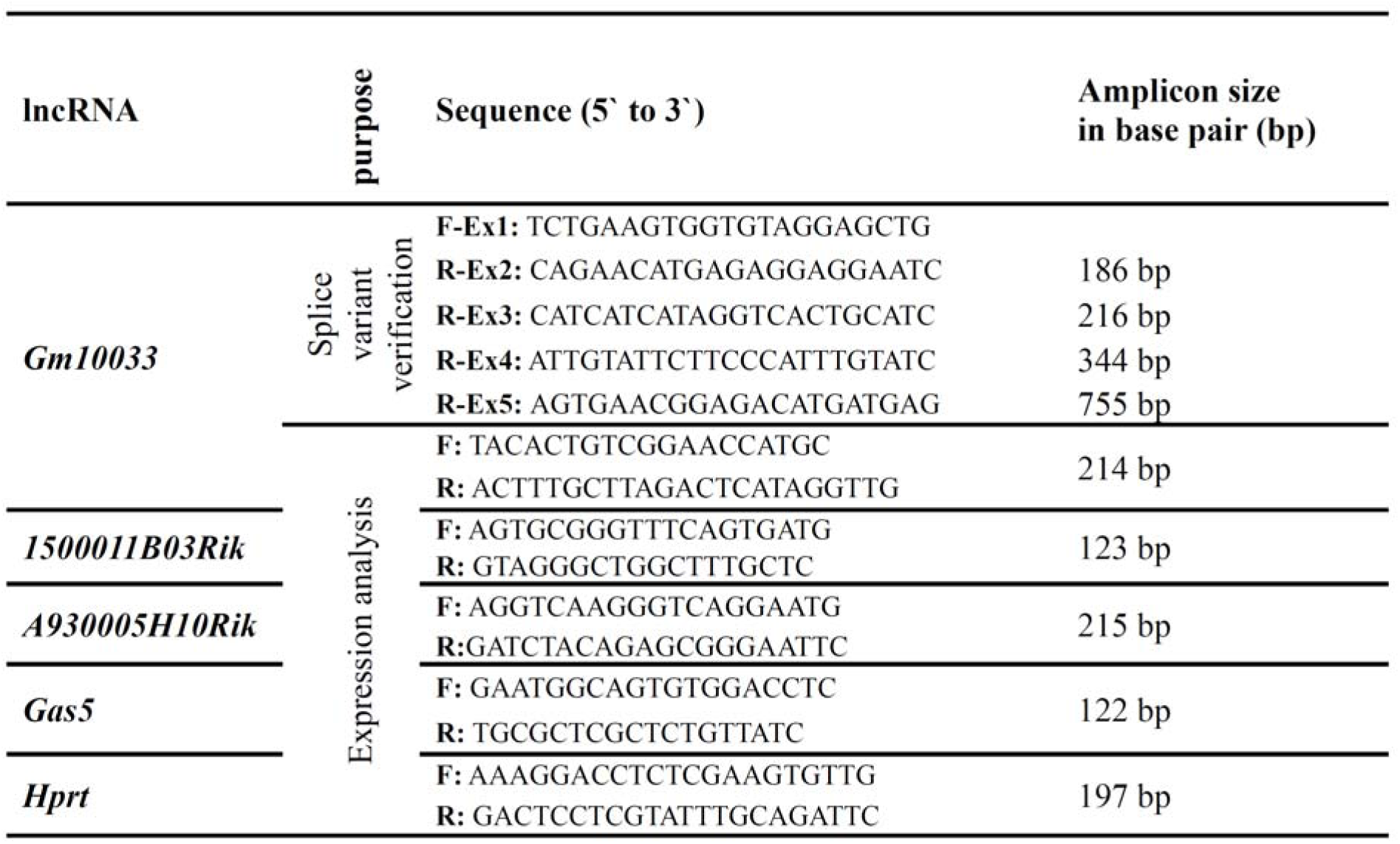
Sequences of RT-PCR primers.

### 2.9. TA cloning

Following amplification, 50 µl of PCR products were run on a 1% agarose gel. PCR fragments were cut out of the gel, purified by the Agarose Gel DNA extraction kit (GeneAll, Korea), ligated into the pTG19-T vector (Sinaclon, Iran), and cloned following the standard protocols. Plasmid DNA was then recovered using the SolGent Plasmid Mini-Prep Kit (SolGent, Korea), sequenced, and analyzed in an Applied Biosystems 3500 sequencing apparatus using M13 universal primers.

### 2.10. Statistical Analysis

The Shapiro-Wilk test was utilized to assess whether the distribution of data were normal. The Student’s t/Mann-Whitney U test was then performed to examine the statistical significances. All analyses and graphs were made using SPSS software, version 16.0 (SPSS Inc., Chicago, USA) and Prism 5.01 (Graph-Pad Prism Software Inc., San Diego, CA). The relative gene expression was calculated according to the 2^-ΔΔCT^ method (39). All data were expressed as mean ± standard error of the mean (SEM) and *p* values of less than 0.05 were defined as statistically significant.

## 3. Results

### 3.1. Three-component regulatory network analysis

To establish a three-component regulatory network, lncRNAs and mRNAs targeted by miRNAs were identified by the above-mentioned methods (section 2.1). The flow chart for network construction is shown in **Figure 1**.

**Fig. 1.**
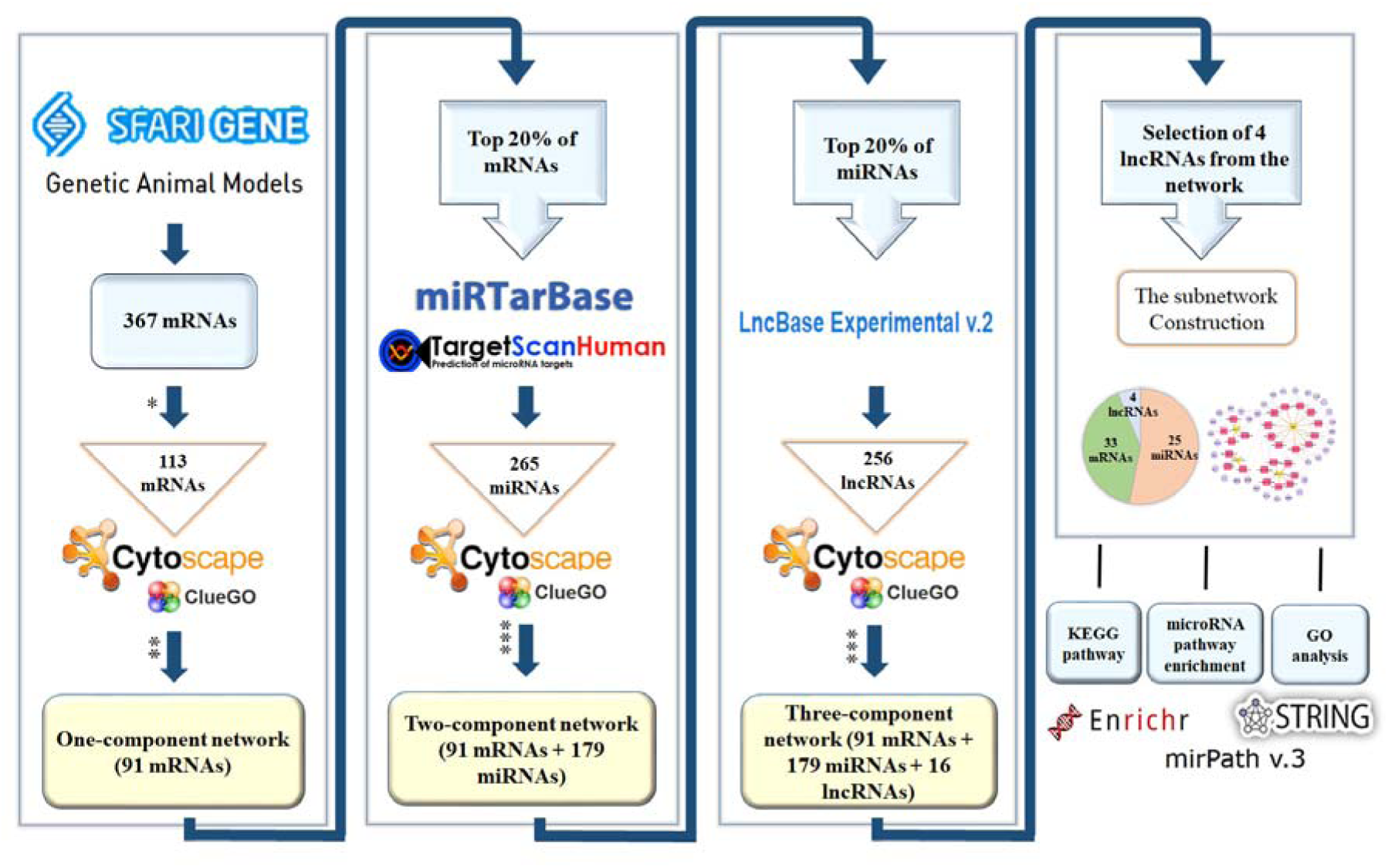
A flowchart of the three-component network construction. The applied filters are represented with stars. One-star (*) shows the filters which were applied on SFARI gene list including a selection of *Mus musculus,* SFARI gene rankings ≤ 4 of evidence strength, and the number of animal model >1. Two-stars (**) show the filters which were applied on CluePedia in Cytoscape to construct a one-component network including the selection of the STRING Action File, activation, inhibition, binding, and post-translational modification with confidence cutoff = 0.5. Three-stars (***) show the filters applied on CluePedia in Cytoscape to construct the two- and three-component networks including the selection of miRanda, miRTarBase, and miRecords databases with a cut off=0.8 as well as filters previously mentioned.

As shown in the **supplementary information 1**, the three-component regulatory network was composed of 91 mRNAs, 179 miRNAs and 16 lncRNAs nodes, as well as 857 edges.

Based on the criteria of high degree centrality, high expression in the mouse frontal lobe and cerebellum (40) and not previously reported in the field of autism, four lncRNAs including *Gm10033*, *1500011B03Rik, A930005H10Rik* and *Gas5* were selected out and a three-component subnetwork was constructed. As shown in Fig. 2, the subnetwork was composed of 33 mRNAs, 25 miRNAs and 4 lncRNAs nodes as well as 94 edges.

**Fig. 2.**
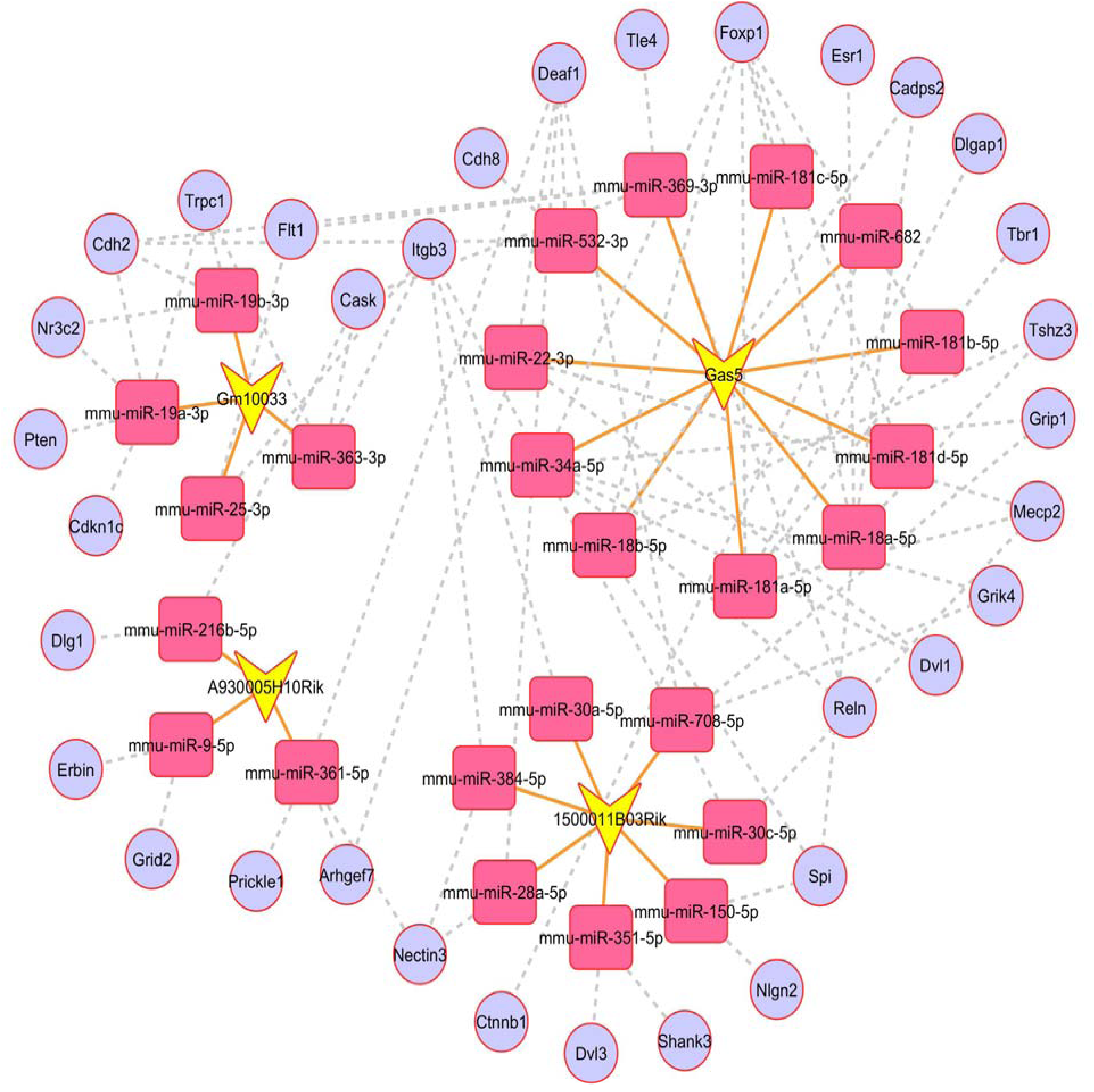
Three-component subnetwork. Light purple circles represent mRNAs, pink rectangles represent miRNAs and yellow V shapes represent lncRNAs. Grey dash and orange lines indicate miRNA-mRNA and miRNA-lncRNA interactions, respectively

### 3.2. Functional implication of subnetwork mRNAs and miRNAs using enrichment analysis

Using KEGG pathway enrichment, we found that mRNAs in the subnetwork were involved in the neuronal differentiation and autism-related pathways like focal adhesion (map04510), glutamatergic synapse (map04724), Wnt signaling pathway (map04310), Hippo signaling pathway (map04390), and PI3K-Akt signaling (map04151) pathway (Fig. 3).

**Fig. 3.**
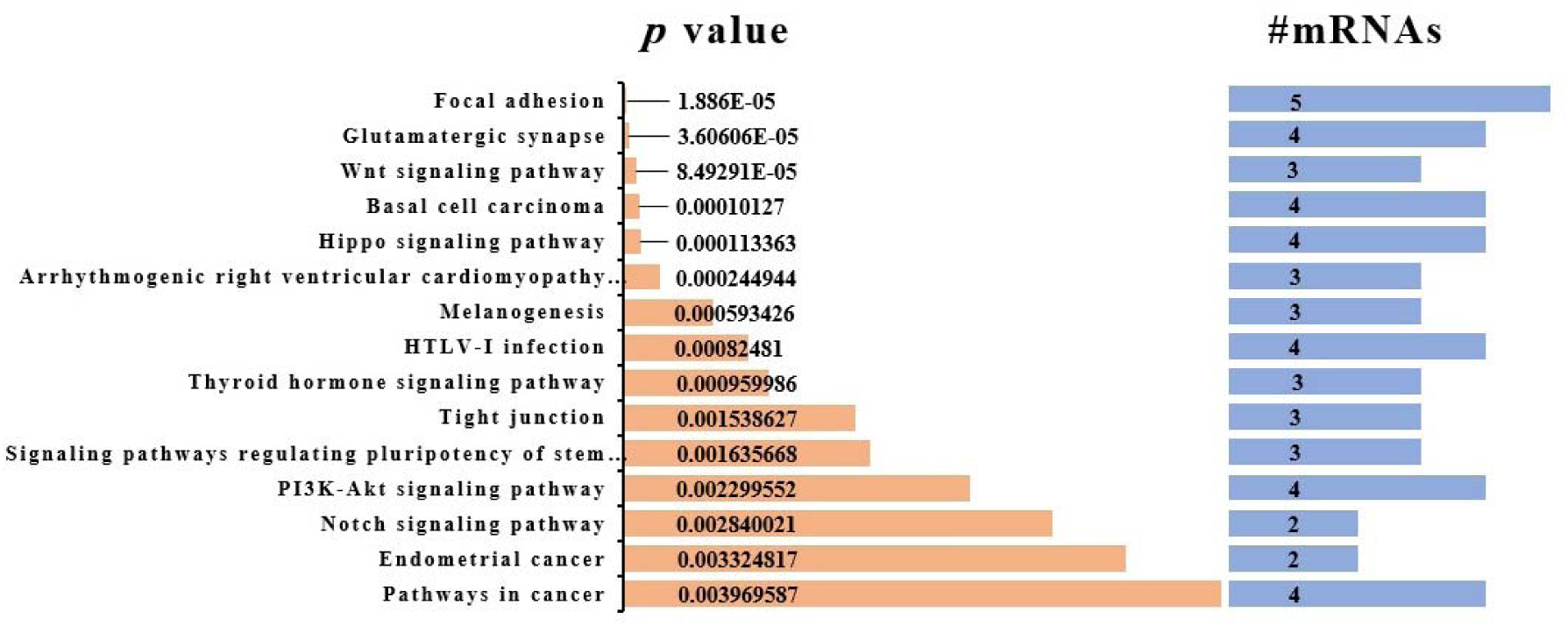
KEGG pathway functional enrichment analysis of mRNAs in the subnetwork. Statistically enriched KEGG pathways utilizing Enrichr (*p* value <0.05) are shown along with the number of mRNAs involved in each pathway

GO analysis revealed that 343 enriched clusters were associated with biological processes and 50 clusters were associated with cellular components (Supplementary Informations 2 and 3). Biological processes such as nervous system development, synapse organization, synaptic transmission, system development, postsynaptic density assembly, synapse assembly, and regulation of nervous system development (Supplementary Information 4A) and cellular components including synapse, postsynaptic membrane, post synapse, synapse part and neuron part (Supplementary Information 4B) were statistically enriched.

In addition, a total of 53 statistically significant KEGG pathways were obtained from the analysis of target genes and involving pathways of all 25 subnetwork miRNAs, using DIANA-mirPath (Supplementary Information 5).

We found that the predicted target genes of all 25 miRNAs in the subnetwork were associated with autism and neurological pathways. These pathways included axon guidance (map04360), MAPK signaling (map04010), TGF beta signaling (map04350), Hippo signaling pathway, neurotrophin signaling pathway (map04722), long-term potentiation (map04720), glutamatergic synapse, and mTOR signaling pathway (map04150) (Supplementary Information 6).

### 3.3. Functional implication of network mRNAs using PPI network analysis

After setting the combined scores greater than 0.9 and hiding the disconnected nodes, the PPI network consisted of 42 proteins. In this network, 19 proteins showed a strong connection (more than 9) with other node proteins (Supplementary Information 7). STRING analysis of these highly-connected proteins showed strong associations with autism-related pathways such as focal adhesion, PI3K-Akt signaling pathway, and neurotrophin signaling pathway (Supplementary Information 8).

### 3.4. Analysis of disease ontology, mammalian phenotype ontology enrichment, and expression profiling of the mouse brain

As a complementary approach, we determined whether our mRNAs in the main network were associated with particular diseases and phenotypes assigned to mouse transgenic and null mice previously developed for these genes. (42). The results of disease enrichment in MGI showed six DO terms including pervasive developmental disorder [DOID:0060040], autism spectrum disorder [DOID:0060041], developmental disorder of mental health [DOID:0060037], disease of mental health [DOID:150], schizophrenia [DOID:5419] and psychotic disorder [DOID:2468].

Furthermore, the mRNAs in the network are involved in the developmental disorder of mental health which is organized into pervasive developmental disorder (PDD) and specific developmental disorder (SDD) as a term with siblings in MGI categories. Autism spectrum disorder, Rett syndrome, and childhood disintegrative disease as a child term are classified in PDD. Furthermore, communication disorder, intellectual disability, and learning disability as a child term are classified in SDD (Fig. 4). Moreover, using mammalian phenotype enrichment in MGI, we found that mRNAs in the network were enriched in 206 statistically significant MP (Supplementary Information 9) including abnormal nervous system physiology (MP:0003633), abnormal social investigation (MP:0001360), abnormal learning/memory/conditioning (MP:0002063), abnormal cognition (MP:0014114) and increased grooming behavior (MP:0001441) which the top 15 statistically enriched MP has been shown in Supplementary Information 10. In fact, concerning the expression data available in MGI, some parts of the brain and central nerves system (CNS) are involved in the above-mentioned MP, including the cranial nerve, tail nervous system, gray matter, ganglion, and brain marginal layer (Supplementary Information 11).

**Fig. 4.**
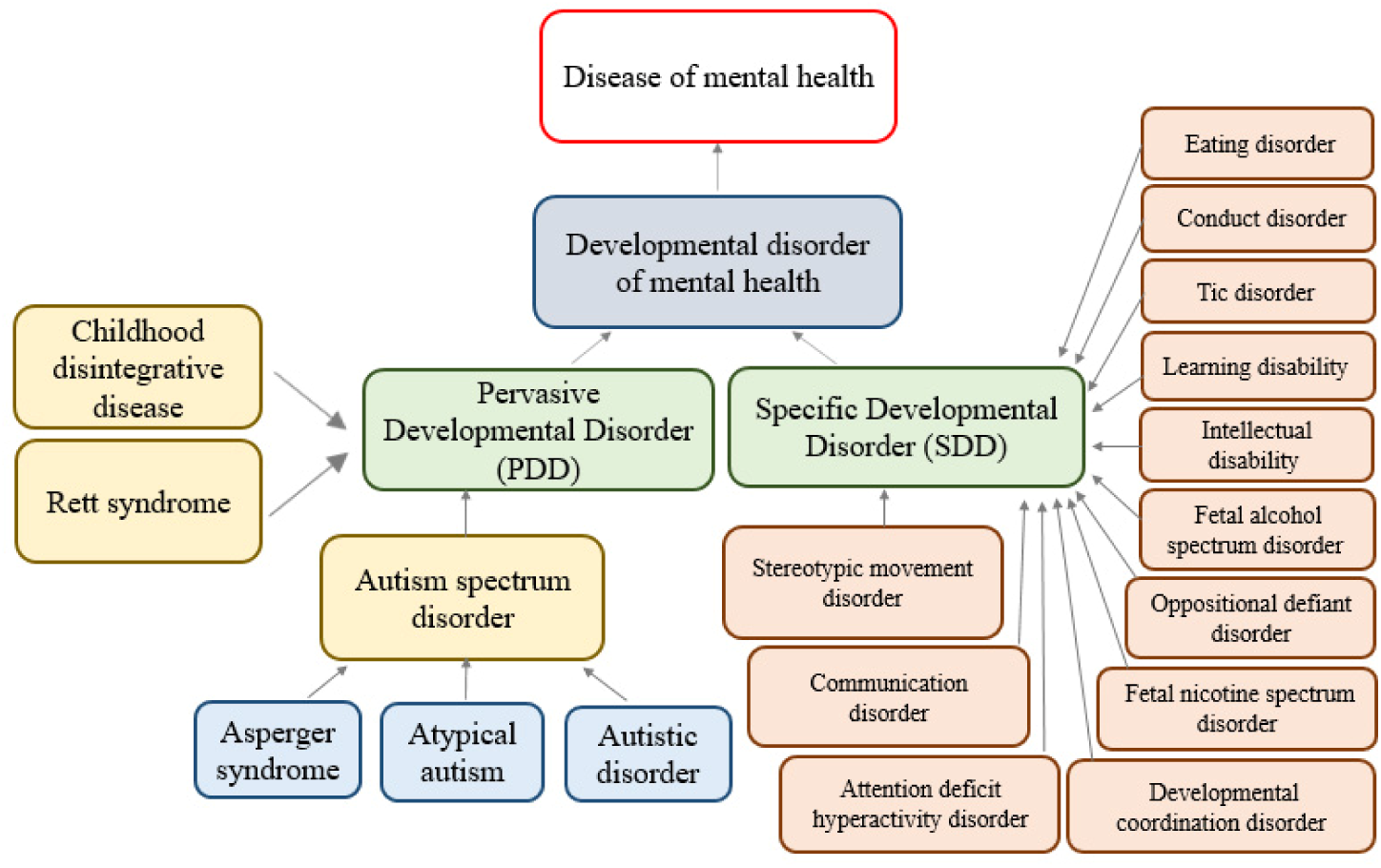
Disease ontology enrichment analysis of mRNAs in the network. The parent term is shown in the blue rectangle, the term with siblings in green, and the child term in other colors

Based on the genome-wide expression data of mouse brain regions samples from the BrainStars project, the expression pattern of the top 6 genes in the main network in the cerebellum (Cb) cortex lobe (Cb lobe), cerebellum nucleus (Cb nucleus), and cerebellar cortex vermis (Cb vermis) which are part of the cerebellum lobe has been shown in Fig. 5. The data indicate *Ctnnb1* has nearly high, *Rhno1*, *Tshz3*, and *Prkdc* have a medium while *Foxp1* and *Grip1* has low expression levels in the cerebellum lobe.

**Fig. 5.**
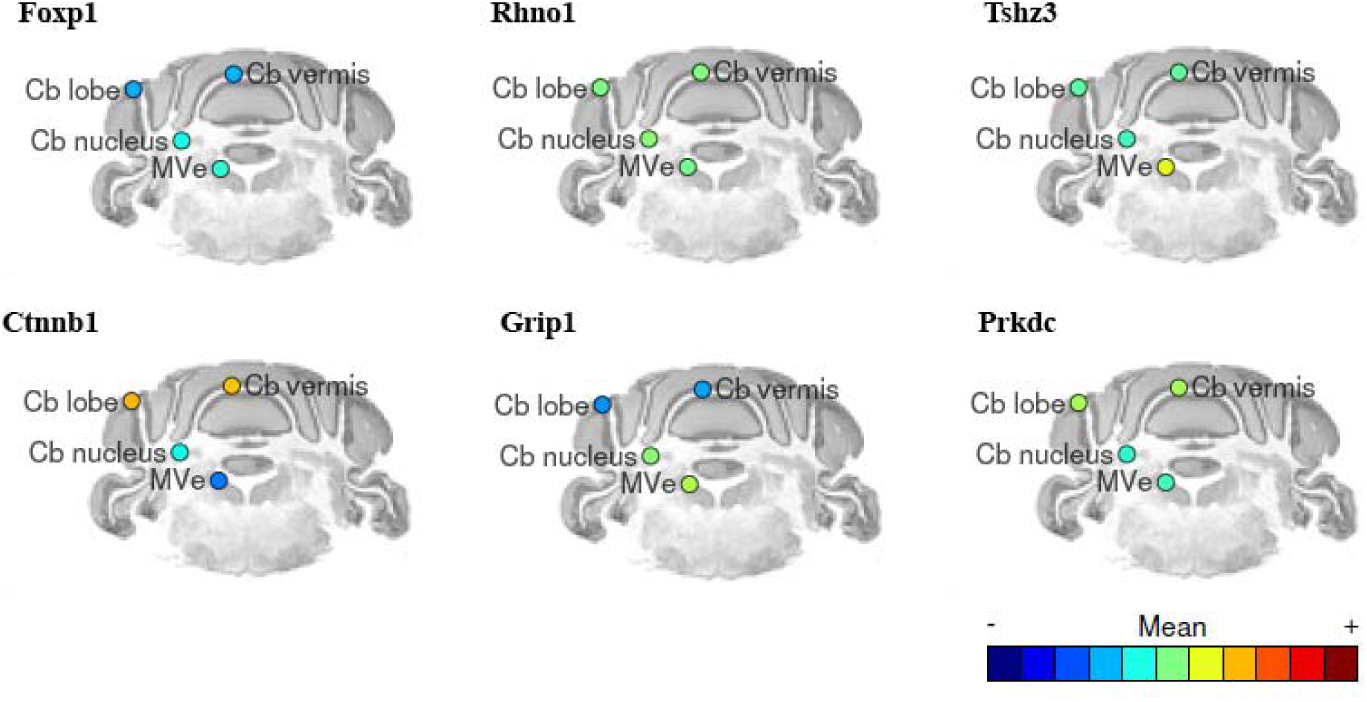
The expression level of the top 6 genes (*Foxp1*, *Rhno1*, *Tshz3*, *Ctnnb1*, *Grip1*, and *Prkdc*) which are sorted based on the degree of centrality in the main network (≥40) in the cerebellum lobe (cerebellum cortex lobe (Cb lobe), cerebellum nucleus (Cb nucleus) and cerebellar cortex vermis (Cb vermis)) of the mouse brain. The level of expression is shown with blue to red from low to the high expression (41)

Considering EMAGE database that provides gene expression patterns during mouse embryogenesis using in situ hybridization, it shows the strong expression of *Ctnnb1* in the midbrain, metencephalon, and telencephalon of mouse embryos between E11.5 to E13 (Supplementary Information 12). Therefore, since the expression of *Ctnnb1* as a part of WNT/β-catenin is necessary for the development of the nervous system, VPA injection in E12.5 can affect brain development and have destructive effects on development.

### 3.5. Analysis of sociability and social novelty

VPA-exposed mice showed less sociability by spending significantly less time in the stranger cage chamber (*p* = 0.0003) compared to the saline group (Fig. 6A). Although the time spent in the empty chamber was not statistically significant between the two groups (Fig. 6B), the thorough analysis showed that VPA-exposed mice spent significantly more time in the central chamber (*p* = 0.0085, data not shown). After introducing a novel mouse into the empty wire cage, VPA-exposed mice tended to stay longer with the stranger mouse (*p* = 0.027; Fig. 6C) than with the novel mouse (*p =* 4.45×10^-6^; Fig. 6D). Moreover, the SI (*p* = 0.005; Fig. 6E) and SPI (*p*=0.002; Fig. 6F) were significantly decreased in the VPA-exposed mice compared to the saline group. Hence, VPA exposure reduced the sociability and social novelty behavior of mice.

**Fig. 6.**
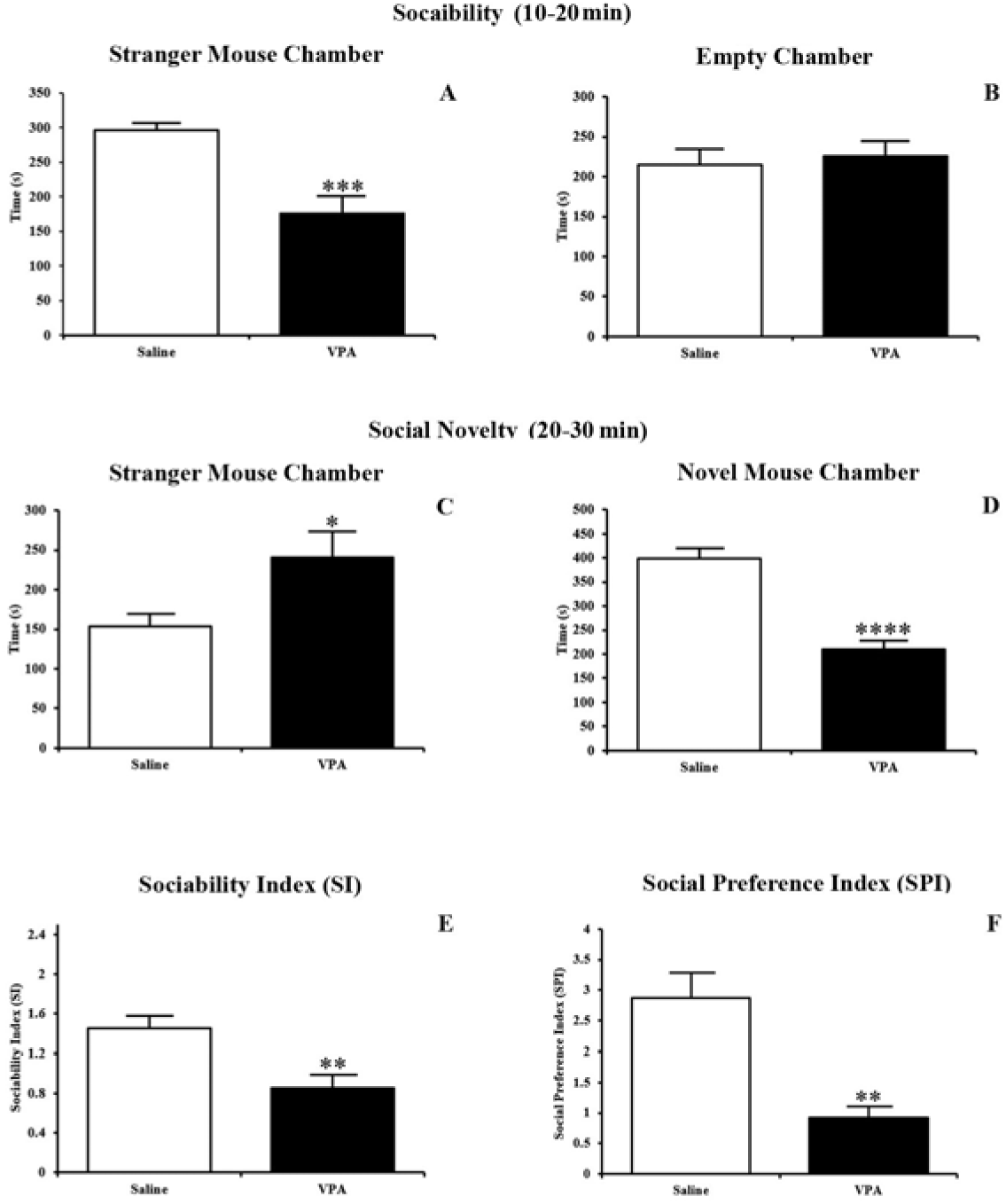
Sociability and social novelty analysis using the three-chamber test. In the second 10 minutes (sociability), the time spent by the subject in the chamber containing the stranger mouse (A) and empty chamber (B) were measured. In the third 10 minutes (social novelty), the time spent by the subject in the chamber containing the stranger mouse (C) and novel mouse chamber (D) were measured. The sociability (E) and social preference indices (F) were equal to time spent in the stranger divided by empty and in the novel divided by the stranger chambers, respectively. Data are expressed as mean±SEM for each category. The stars show statistically significant differences between the VPA-exposed mice and saline group (*p < 0.05, **p < 0.01 and ***p < 0.001)

### 3.6. Analysis of locomotor activity, self-grooming and anxiety-like behaviors

VPA-exposed mice showed significantly higher (*p =* 0.001) locomotor activity when compared to the saline group (Fig. 7A). In addition, the VPA-exposed mice groomed significantly more (*p* = 0.006) than the saline group, as a sign of repetitive behavior (Fig. 7B). Furthermore, as a sign of anxiety-like behaviors, the time spent in the outer and inner zones by VPA-exposed mice was significantly more (*p* = 0.009) (Fig. 7C) and less (*p* = 0.0005) (Fig. 7D) than the saline group, respectively. These findings indicated an increase in locomotor activity, self-grooming, and anxiety-like behaviors in VPA-exposed mice.

**Fig. 7.**
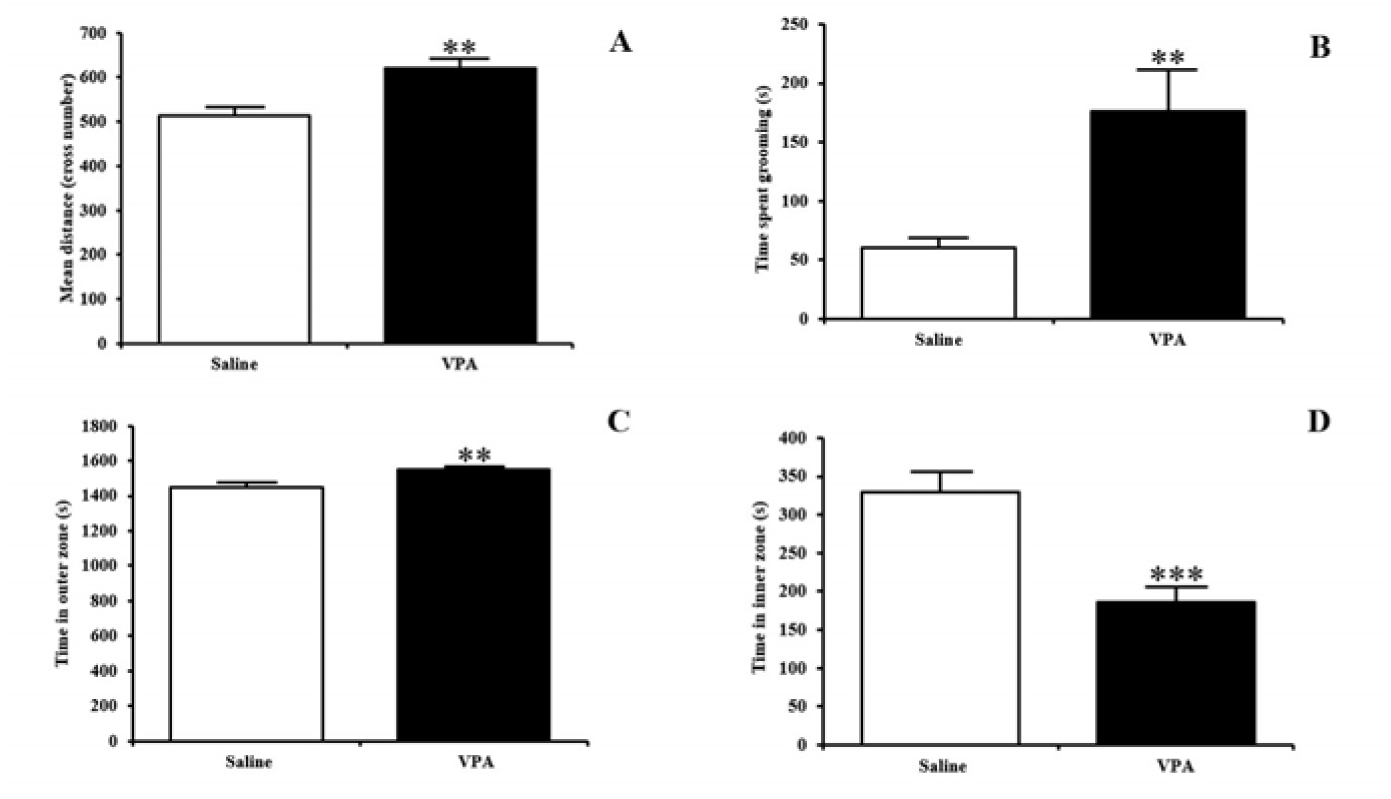
Analysis of locomotor activity, self-grooming, and anxiety-like behaviors using open field test. The total locomotor activity was calculated through crossing one of the squares with all four paws (A), self-grooming (as a sign of repetitive behavior) (B), the time spent in the outer (C) and inner (D) zones is shown as a sign of anxiety-like behaviors. Data are expressed as mean±SEM. The stars show statistically significant differences between the VPA-exposed mice and saline group (**p < 0.01 and ***p < 0.001)

### 3.7. Analysis of long-term memory via passive avoidance testing

VPA-exposed mice showed significantly reduced latency in the retention trial (*p* = 0.001), when compared to the saline group (Fig. 8), confirming impairment of their long-term memory.

**Fig. 8.**
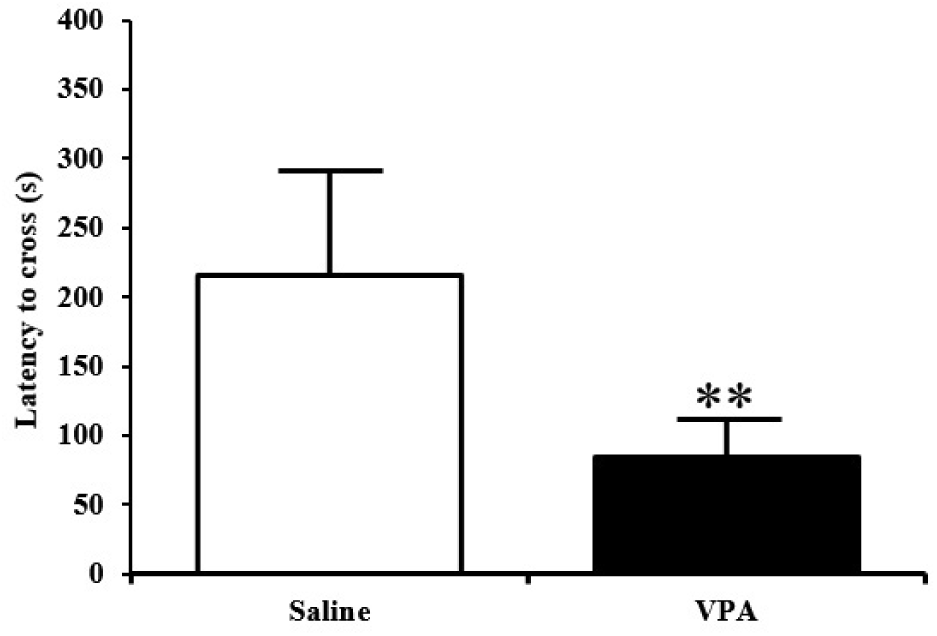
Analysis of long-term memory using passive avoidance test. The latency of the subject to escape into the dark chamber was measured in two consecutive days (acquisition and retention trials). Data are expressed as mean±SEM. The stars show statistically significant differences between the VPA-exposed mice and saline group (**p < 0.01)

### 3.8. PCR optimization and identification of Gm10033-**Δ**Ex2 as a novel splice variant of the *Gm10033*

To specifically amplify the four candidate lncRNAs, optimization was carried out by conventional RT-PCR reaction. Agarose gel electrophoresis of the PCR products displayed specific bands with the expected sizes for *1500011B03Rik, A930005H10Rik,* and *Gas5* (Supplementary Information 13).

For *Gm10033*, we expect a specific band with a size of 216 bp. However, conventional RT-PCR using forward and reverse primers (designed on exon 1 and 3, respectively) resulted in an extra band with an approximate size of 110 bp (data not shown). We accordingly gel-extracted the two bands, cloned them into a TA vector, and bi-directionally sequenced them. The sequencing results revealed the existence of a novel shorter splice variant, *Gm10033-*Δ*Ex2* with the removal of exon 2 (110 bp) (Fig. 9).

**Fig. 9.**
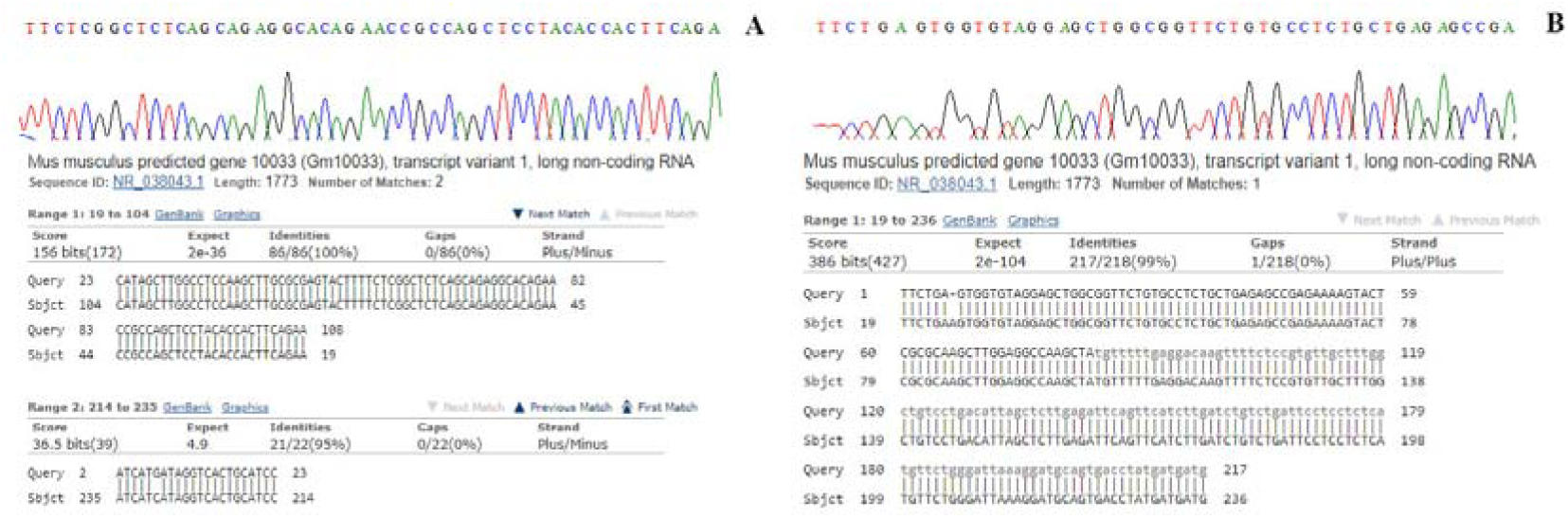
DNA sequencing analysis of two bands following *Gm10033* RT-PCR amplification with F-Ex1 and R-Ex3 primers. A part of a sequence electropherogram of the novel shorter splice variant, *Gm10033-*Δ*Ex2* (A) and *Gm10033* transcript (B) and their nucleotide BLAST are represented

For further investigation of *Gm10033-*Δ*Ex2,* three extra reverse primers located on exons 2, 4, and 5 were designed (Supplementary Information 14).

As expected, in addition to the specific bands with predicted sizes for each pair of primers (corresponding to *Gm10033* (NR_038043.2)), one more band resulting from amplification of *Gm10033-*Δ*Ex2* was observed (Supplementary Information 15).

The tissue distribution of *Gm10033-*Δ*Ex2* expression was then characterized in a panel of mouse adult tissues by RT-PCR. Except for the gastrocnemius muscle, *Gm10033-*Δ*Ex2* was detected in the liver, kidney, heart, lung, testis, and the whole brain (Supplementary Information 16).

Altogether, these results suggest the existence and tissue-specific expression pattern of a novel shorter splice variant for *Gm10033*. A specific band with the expected size of 214 bp (first lane in Supplementary Information 13) was obtained by RT-PCR amplification of *Gm10033* with a new pair of primers designed on exon 5 which were subsequently used to assess the gene expression. To confirm the specific amplification of the four candidate lncRNAs, PCR products were sequenced and database comparisons using nucleotide BLAST revealed a 100 % identity of the amplified PCR products with the corresponding transcripts (Supplementary Information 17).

### 3.9. Quantitative validation of the four candidate lncRNAs in mouse brain

Compared to the saline group, a statistically significant increased expression of *Gm10033* was observed in the frontal lobe and cerebellum of VPA-exposed mice (*p* = 0.011 and *p* = 0.001, respectively) (Fig. 10A-B). Significant alteration of *1500011B03Rik* with a 1.5-fold increase was observed in the frontal lobe of VPA-exposed mice (*p* = 0.040). In the cerebellum, a more significant change of *1500011B03Rik* expression with an 8.55-fold increase was demonstrated (*p* = 0.0001) (Fig. 10C-D). VPA-exposed mice exhibited an increased expression of *A930005H10Rik* in the frontal lobe (*p* = 0.054). The same trend, although statistically significant, was likewise observed for the *A930005H10Rik* expression level in the cerebellum of VPA-exposed mice (*p* = 0.018) (Fig. 10E-F). The expression of *Gas5* was statistically increased in the frontal lobe and cerebellum of VPA-exposed mice (*p* = 0.003) (Fig. 10G-H). Taken together, differential expression of *Gm10033, 1500011B03Rik, A930005H10Rik,* and *Gas5* in the brain of VPA-exposed mice introduced them as potentially novel autism-associated lncRNAs.

**Fig. 10.**
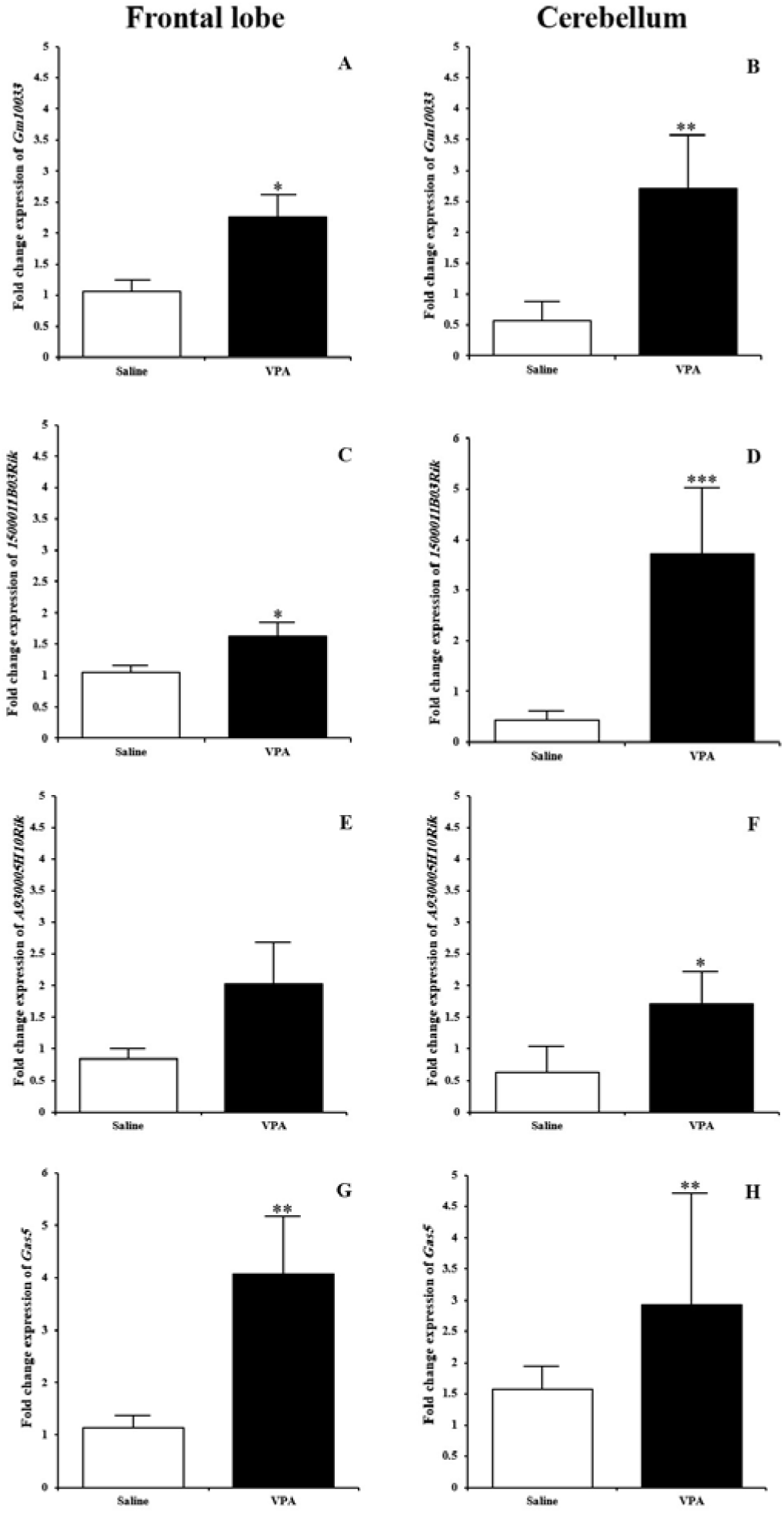
Relative expression of four candidate lncRNAs using quantitative real time RT-PCR. The left panels represent the relative expression of *Gm10033* (A), *1500011B03Rik* (C), *A930005H10Rik* (E), and *Gas5* (G) in the frontal lobe of VPA-exposed mice compared to the saline group. The right panels represent the relative expression of *Gm10033* (B), *1500011B03Rik* (D), *A930005H10Rik* (F), and *Gas5* (H) in the cerebellum of VPA-exposed mice compared to the saline group. Data are expressed as mean±SEM. The stars show statistically significant differences (*p < 0.05, **p < 0.01 and ***p < 0.001)

## 4. Discussion

Although there are multiple studies describing networks of mRNAs, miRNAs, and lncRNAs to find autism-associated non-coding genes, these characterized genes are only the tip of an iceberg, as most of the non-coding RNAs, including lncRNAs, have not been investigated so far.

Based on SFARI data, we constructed an autism-specific mRNA-miRNA-lncRNA network revealing a complex regulatory relationship between different types of genes. Using a bioinformatics approach, four candidate lncRNAs, comprising *Gm10033*, *1500011B03Rik*, *A930005H10Rik,* and *Gas5,* were screened out for further analysis. So far, there have been no detailed reports on studying these lncRNAs in the context of autism. KEGG pathway and GO analysis indicated that mRNAs and miRNAs interacted with the candidate lncRNAs mainly function in synapse organization, synaptic transmission, synapse assembly, and regulation of nervous system development. Based on KEGG pathway analysis, several pathways were enriched, including glutamatergic synapse, Wnt signaling pathway, neurotrophin signaling pathway, Hippo signaling pathway, and PI3K-Akt signaling pathway which were previously reported as dysregulated in autism (42–45). Furthermore, concerning the disease ontology enrichment, we revealed the association between mRNAs in the network with PDD and SDD. According to the previous studies, PDD refers to the group of heterogeneous conditions that formed a continuum disability with sharing a core triad symptom including qualitative impairments in reciprocal social interactions, verbal and nonverbal communication, and repetitive and stereotyped patterns of interests and behavior which autism is classified as a subset to these group of disorders (46, 47). Moreover, several symptoms that are seen in patients or mouse models of autism including delays in developmental domains such as motor function, language and speech development, and uncanny intelligence or poor educational conditions, are coordinated to SDD definition (48). Therefore, these candidates’ lncRNAs might play important and functional roles by interacting with key mRNA(s) and miRNA(s) that in turn may contribute to autism.

Exposure to valproic acid, a widely used anticonvulsant drug, during pregnancy has been associated with an increased incidence of autism (49). VPA-exposed rodents showed the core behavioral signs of autism as well as molecular changes associated with the disorder (50). In the current study, reduced sociability and social novelty, increased locomotor activity, self-grooming, anxiety-like behaviors, and impairment of long-term memory in VPA-exposed mice confirmed the validity of the animal model.

Then, we speculated that the four candidates lncRNAs initially screened out from a three-component regulatory network may show a link with autism because our enrichment analysis predicted that these lncRNAs and their associated mRNAs and miRNAs are involved in multiple autism and neurological pathways. Therefore, we performed a quantitative real time RT-PCR to examine the expression of these four candidate lncRNAs in the frontal lobe and cerebellum of VPA-exposed mice. Increased expression of *Gm10033*, *1500011B03Rik*, *A930005H10Rik,* and *Gas5* were observed in both frontal lobe and cerebral brain samples of VPA-exposed mice. Although differential expression of these lncRNAs was previously reported in other physiological and pathological states (47, 48, 51), there are no reports of involving these lncRNAs in autism or other neurological disorders. However, decreased expression of *Gas5* was reported in the whole brain tissue of mouse embryos treated with VPA on GD 8, as a chemically induced neural tube defects (NTDs) mouse model (52). As several differences exist in our experimental design with theirs, such as VPA dose, day of VPA injection, and the tissue investigated, the results cannot simply be compared between these two studies.

Moreover, in the present study, we identified a novel splice variant of *Gm10033*, designated as *Gm10033-*Δ*Ex2*. Expression of *Gm10033-*Δ*Ex2* was observed in various mouse tissues, including liver, kidney, heart, lung, testis, and brain. The presence of distinct variants of lncRNA with unknown functions, *Gm10033*, suggests that it may play a more complex regulatory role as lncRNA in a wide range of biological processes.

## 5. Conclusion

Utilizing an autism-specific three-component network comprising mRNAs, lncRNAs, and miRNAs, we have identified a list of four candidate lncRNAs. These lncRNAs and their associated mRNAs and miRNAs were found to be associated with neurologically relevant pathways and functions. Experimental validation of *Gm10033*, *1500011B03Rik*, *A930005H10Rik,* and *Gas5* in the brain of VPA-exposed mice introduces them as potentially novel autism-associated lncRNAs. Further studies are required to unravel the functional role of these lncRNAs in the pathobiology of autism. The present work suggests opportunities to define subgroups of patients within a large heterogeneous clinical category and to examine common treatment targets across distinct neurodevelopmental trajectories (53).

## Authors’ contributions

MG carried out the experiments. MG and PN performed bioinformatics analyses. MEB, MM and PN designed and directed the project. MG and PN wrote the manuscript with input from all authors. All authors have seen and agreed to the final submitted version.

## Ethics approval statement

All animal experiments were approved by the institutional animal ethics review board of Isfahan University of Medical Sciences

## Abbreviations

Cb: Cerebellum
CDC: Centers for Disease Control and Prevention
cDNA: Complementary DNA
CNS: Central Nerves System
DO: Disease Ontology
FDR: False Discovery Rate
GD: Gestational Day
GO: Gene Ontology
KEGG: Kyoto Encyclopedia of Genes and Genomes
lncRNA: long noncoding RNA
miRNA: microRNA
MP: Mammalian Phenotype
MGI: Mouse Genome Informatics
NTD: Neural Tube Defect
PCR: Polymerase Chain Reaction
PDD: Pervasive Developmental Disorder
PPI: Protein-Protein Interaction
RT-PCR: Reverse Transcription Polymerase Chain Reaction
SDD: Specific Developmental Disorder
SI: Sociability Index
SFARI: Simons Foundation Autism Research Initiative
SPI: Social Preference Index
VPA: Valproic Acid

## Acknowledgements

The authors would like to thank Farnaz Mohammadtaheri, Nahid Mirmohammadsadeghi and Kobra Moradzadeh for their help in experimental and bioinformatic analysis.

## Supplementary Information legends

**Supplementary Information 1** The three-component network. Light purple circles represent mRNAs, pink rectangles represent miRNAs and yellow V shapes represent lncRNAs. Grey dash and orange lines indicate miRNA-mRNA and miRNA-lncRNA interactions, respectively

**Supplementary Information 2** GO biological processes associated with the mRNAs in the network

**Supplementary Information 3** GO cellular components associated with the mRNAs in the network

**Supplementary Information 4** Gene ontology enrichment analysis of mRNAs in the subnetwork. Top 15 statistically enriched biological processes (A) and cellular components (B) identified by employing STRING. False discovery rate (FDR) and number of mRNAs involving in each pathway are represented

**Supplementary Information 5** Retrieved KEGG pathways from the analysis of miRNAs using DIANA-mirPath

**Supplementary Information 6** Pathway analysis of miRNAs in the subnetwork. Top 25 statistically enriched pathways utilizing DIANA-mirPath (*p* value <0.05) are shown along with the number of mRNAs involving in each pathway

**Supplementary Information 7** STRING database analysis of the PPI network of mRNAs in the main network. A total of 42 mRNAs were remained with the combined scores greater than 0.9 in the STRING network. The numbers of connections with other node proteins are represented in each bar (A) as well as an overview of the PPI network (B)

**Supplementary Information 8** PPI network analysis of the mRNAs in the network

**Supplementary Information 9** Mammalian phenotype enrichment using MGI database

**Supplementary Information 10** Mammalian phenotype ontology enrichment analysis of mRNAs in the network. Considering MGI, top 15 statistically enriched phenotypes (p value <0.05) are shown based on the importance of each pathway

**Supplementary Information 11** The part of the mouse brain and CNS involved in some mammalian phenotypes retrieved from MGI database

**Supplementary Information 12** Expression pattern of *Ctnnb1* during mouse embryogenesis. The expression levels in each part of the brain of an embryo from E11.5 to E13 are shown with colors from red (high expression) to pale blue (low expression) (54, 55)

**Supplementary Information 13** Optimization of PCR conditions for four selected lncRNAs. Agarose gel electrophoresis of *Gm10033*, *1500011B03Rik*, *A930005H10Rik* and *Gas5* showed specific bands with the expected sizes. The first lane represents a 100 bp DNA ladder

**Supplementary Information 14** Schematic representation of exon organization in the *Gm10033* transcript (NR_038043.2). The boxes represent exons and the diagonal pattern in the exon 2 represents the 110 bp which is lacking in the novel shorter splice variant, Gm10033-ΔEx2. The arrows represent positions of RT-PCR primer

**Supplementary Information 15** Whole length analysis of *Gm10033* (NR_038043.2) and Gm10033-ΔEx2 transcripts using conventional RT-PCR. The designed forward primer for exon 1 (F-Ex1) was used with all four reverse primers (R-Ex2-5). A single band in the second lane resulted from RT-PCR amplification of Gm10033 transcript having the exon 2. The first lane represents a 100 bp DNA ladder

**Supplementary Information 16** The tissue distribution of Gm10033-ΔEx2 expression in a panel of mouse adult tissues. Total RNA extracted from liver, kidney, heart, lung, testis, gastrocnemius muscle and whole brain were used as a template in RT-PCR reaction using F-Ex1 and R-Ex3 primers. The first lane represents a 100 bp DNA ladder

**Supplementary Information 17** DNA sequencing analysis of four candidate lncRNAs. A part of a sequence electropherogram of *Gm10033* (A*) 1500011B03Rik* (B), *A930005H10Rik* (C) and *Gas5* (D) and their nucleotide BLAST are represented

## References

1. Halfon N, Kuo AA. What DSM-5 could mean to children with autism and their families. JAMA Pediatr. 2013;167(7):608–13.

2. Abu-Elneel K, Liu T, Gazzaniga FS, Nishimura Y, Wall DP, Geschwind DH, et al. Heterogeneous dysregulation of microRNAs across the autism spectrum. Neurogenetics. 2008;9(3):153–61.

3. Grzadzinski R, Huerta M, Lord C. DSM-5 and autism spectrum disorders (ASDs): an opportunity for identifying ASD subtypes. Mol Autism. 2013;4(1):12.

4. Halfon N, Kuo AA. What DSM-5 could mean to children with autism and their families. JAMA pediatrics. 2013;167(7):608–13.

5. Baio J, Wiggins L, Christensen DL, Maenner MJ, Daniels J, Warren Z, et al. Prevalence of Autism Spectrum Disorder Among Children Aged 8 Years - Autism and Developmental Disabilities Monitoring Network, 11 Sites, United States, 2014. MMWR Surveill Summ. 2018;67(6):1–23.

6. Ander BP, Barger N, Stamova B, Sharp FR, Schumann CM. Atypical miRNA expression in temporal cortex associated with dysregulation of immune, cell cycle, and other pathways in autism spectrum disorders. Mol Autism. 2015;6:37.

7. Huang F, Long Z, Chen Z, Li J, Hu Z, Qiu R, et al. Investigation of Gene Regulatory Networks Associated with Autism Spectrum Disorder Based on MiRNA Expression in China. PLoS One. 2015;10(6):e0129052.

8. Gregg JP, Lit L, Baron CA, Hertz-Picciotto I, Walker W, Davis RA, et al. Gene expression changes in children with autism. Genomics. 2008;91(1):22–9.

9. Tang J, Yu Y, Yang W. Long noncoding RNA and its contribution to autism spectrum disorders. CNS Neurosci Ther. 2017;23(8):645–56.

10. Pastori C, Wahlestedt C. Involvement of long noncoding RNAs in diseases affecting the central nervous system. RNA Biol. 2012;9(6):860–70.

11. Voineagu I, Wang X, Johnston P, Lowe JK, Tian Y, Horvath S, et al. Transcriptomic analysis of autistic brain reveals convergent molecular pathology. Nature. 2011;474(7351):380–4.

12. Ziats MN, Rennert OM. Aberrant expression of long noncoding RNAs in autistic brain. J Mol Neurosci. 2013;49(3):589–93.

13. Wang Y, Zhao X, Ju W, Flory M, Zhong J, Jiang S, et al. Genome-wide differential expression of synaptic long noncoding RNAs in autism spectrum disorder. Transl Psychiatry. 2015;5:e660.

14. Banerjee-Basu S, Packer A. SFARI Gene: an evolving database for the autism research community. Dis Model Mech. 2010;3(3-4):133–5.

15. Bindea G, Galon J, Mlecnik B. CluePedia Cytoscape plugin: pathway insights using integrated experimental and in silico data. Bioinformatics. 2013;29(5):661–3.

16. Shannon P, Markiel A, Ozier O, Baliga NS, Wang JT, Ramage D, et al. Cytoscape: a software environment for integrated models of biomolecular interaction networks. Genome Res. 2003;13(11):2498–504.

17. Chou CH, Shrestha S, Yang CD, Chang NW, Lin YL, Liao KW, et al. miRTarBase update 2018: a resource for experimentally validated microRNA-target interactions. Nucleic Acids Res. 2018;46(D1):D296–D302.

18. Agarwal V, Bell GW, Nam JW, Bartel DP. Predicting effective microRNA target sites in mammalian mRNAs. Elife. 2015;4.

19. Paraskevopoulou MD, Vlachos IS, Karagkouni D, Georgakilas G, Kanellos I, Vergoulis T, et al. DIANA-LncBase v2: indexing microRNA targets on non-coding transcripts. Nucleic Acids Res. 2016;44(D1):D231–8.

20. Kuleshov MV, Jones MR, Rouillard AD, Fernandez NF, Duan Q, Wang Z, et al. Enrichr: a comprehensive gene set enrichment analysis web server 2016 update. 2016;44(W1):W90–W7.

21. Vlachos IS, Kostoulas N, Vergoulis T, Georgakilas G, Reczko M, Maragkakis M, et al. DIANA miRPath v.2.0: investigating the combinatorial effect of microRNAs in pathways. Nucleic Acids Res. 2012;40(Web Server issue):W498–504.

22. Szklarczyk D, Franceschini A, Wyder S, Forslund K, Heller D, Huerta-Cepas J, et al. STRING v10: protein-protein interaction networks, integrated over the tree of life. Nucleic Acids Res. 2015;43(Database issue):D447–52.

23. The Gene Ontology C. The Gene Ontology Resource: 20 years and still GOing strong. Nucleic Acids Res. 2019;47(D1):D330–D8.

24. Eppig JT. Mouse Genome Informatics (MGI) Resource: Genetic, Genomic, and Biological Knowledgebase for the Laboratory Mouse. ILAR J. 2017;58(1):17–41.

25. Bello SM, Smith CL, Eppig JT. Allele, phenotype and disease data at Mouse Genome Informatics: improving access and analysis. Mamm Genome. 2015;26(7-8):285–94.

26. Kataoka S, Takuma K, Hara Y, Maeda Y, Ago Y, Matsuda T. Autism-like behaviours with transient histone hyperacetylation in mice treated prenatally with valproic acid. Int J Neuropsychopharmacol. 2013;16(1):91–103.

27. Schneider T, Roman A, Basta-Kaim A, Kubera M, Budziszewska B, Schneider K, et al. Gender-specific behavioral and immunological alterations in an animal model of autism induced by prenatal exposure to valproic acid. Psychoneuroendocrinology. 2008;33(6):728–40.

28. Yang EJ, Ahn S, Lee K, Mahmood U, Kim HS. Early Behavioral Abnormalities and Perinatal Alterations of PTEN/AKT Pathway in Valproic Acid Autism Model Mice. PLoS One. 2016;11(4):e0153298.

29. Kaidanovich-Beilin O, Lipina T, Vukobradovic I, Roder J, Woodgett JR. Assessment of social interaction behaviors. J Vis Exp. 2011 (48).

30. Chau DK, Choi AY, Yang W, Leung WN, Chan CW. Downregulation of glutamatergic and GABAergic proteins in valproric acid associated social impairment during adolescence in mice. Behav Brain Res. 2017;316:255–60.

31. Choi CS, Gonzales EL, Kim KC, Yang SM, Kim JW, Mabunga DF, et al. The transgenerational inheritance of autism-like phenotypes in mice exposed to valproic acid during pregnancy. Sci Rep. 2016;6:36250.

32. Wei R, Li Q, Lam S, Leung J, Cheung C, Zhang X, et al. A single low dose of valproic acid in late prenatal life alters postnatal behavior and glutamic acid decarboxylase levels in the mouse. Behav Brain Res. 2016;314:190–8.

33. Olexova L, Stefanik P, Krskova L. Increased anxiety-like behaviour and altered GABAergic system in the amygdala and cerebellum of VPA rats - An animal model of autism. Neurosci Lett. 2016;629:9–14.

34. Dachtler J, Ivorra JL, Rowland TE, Lever C, Rodgers RJ, Clapcote SJ. Heterozygous deletion of alpha-neurexin I or alpha-neurexin II results in behaviors relevant to autism and schizophrenia. Behav Neurosci. 2015;129(6):765–76.

35. Soeda F, Hirakawa E, Inoue M, Shirasaki T, Takahama K. Cloperastine rescues impairment of passive avoidance response in mice prenatally exposed to diethylstilbestrol. Environ Toxicol. 2014;29(2):216–25.

36. Brown SL, Kolozsvary A, Liu J, Ryu S, Kim JH. Histone deacetylase inhibitors protect against and mitigate the lethality of total-body irradiation in mice. Radiat Res. 2008;169(4):474–8.

37. Zhou J, Zhang X, Ren J, Wang P, Zhang J, Wei Z, et al. Validation of reference genes for quantitative real-time PCR in valproic acid rat models of autism. Mol Biol Rep. 2016;43(8):837–47.

38. Hall TA, editor BioEdit: a user-friendly biological sequence alignment editor and analysis program for Windows 95/98/NT. Nucleic acids symposium series; 1999: [London]: Information Retrieval Ltd., c1979–c2000.

39. Livak KJ, Schmittgen TD. Analysis of relative gene expression data using real-time quantitative PCR and the 2(-Delta Delta C(T)) Method. Methods. 2001;25(4):402–8.

40. Yue F, Cheng Y, Breschi A, Vierstra J, Wu W, Ryba T, et al. A comparative encyclopedia of DNA elements in the mouse genome. Nature. 2014;515(7527):355–64.

41. Kasukawa T, Masumoto KH, Nikaido I, Nagano M, Uno KD, Tsujino K, et al. Quantitative expression profile of distinct functional regions in the adult mouse brain. PLoS One. 2011;6(8):e23228.

42. Zhang Y, Yuan X, Wang Z, Li R. The canonical Wnt signaling pathway in autism. CNS Neurol Disord Drug Targets. 2014;13(5):765–70.

43. Nickl-Jockschat T, Michel TM. The role of neurotrophic factors in autism. Mol Psychiatry. 2011;16(5):478–90.

44. Zhou J, Parada LF. PTEN signaling in autism spectrum disorders. Curr Opin Neurobiol. 2012;22(5):873–9.

45. Gazestani VH, Pramparo T, Nalabolu S, Kellman BP, Murray S, Lopez L, et al. A perturbed gene network containing PI3K/AKT, RAS/ERK, WNT/β-catenin pathways in leukocytes is linked to ASD genetics and symptom severity. Nature Neuroscience. 2019;22:1624–34.

46. Fombonne E. Epidemiology of pervasive developmental disorders. Pediatr Res. 2009;65(6):591–8.

47. Mourtada-Maarabouni M, Pickard MR, Hedge VL, Farzaneh F, Williams GT. GAS5, a non-protein-coding RNA, controls apoptosis and is downregulated in breast cancer. Oncogene. 2009;28(2):195–208.

48. Mozhui K, Karlsson RM, Kash TL, Ihne J, Norcross M, Patel S, et al. Strain differences in stress responsivity are associated with divergent amygdala gene expression and glutamate-mediated neuronal excitability. J Neurosci. 2010;30(15):5357–67.

49. Christensen J, Gronborg TK, Sorensen MJ, Schendel D, Parner ET, Pedersen LH, et al. Prenatal valproate exposure and risk of autism spectrum disorders and childhood autism. JAMA. 2013;309(16):1696–703.

50. Schneider T, Przewlocki R. Behavioral alterations in rats prenatally exposed to valproic acid: animal model of autism. Neuropsychopharmacology. 2005;30(1):80–9.

51. MacLaren EJ, Sikela JM. Cerebellar gene expression profiling and eQTL analysis in inbred mouse strains selected for ethanol sensitivity. Alcohol Clin Exp Res. 2005;29(9):1568–79.

52. Okada A, Kushima K, Aoki Y, Bialer M, Fujiwara M. Identification of early-responsive genes correlated to valproic acid-induced neural tube defects in mice. Birth Defects Res A Clin Mol Teratol. 2005;73(4):229–38.

53. Jacob S, Wolff JJ, Steinbach MS, Doyle CB, Kumar V, Elison JT. Neurodevelopmental heterogeneity and computational approaches for understanding autism. Transl Psychiatry. 2019;9(1):63.

54. Mucenski ML, Wert SE, Nation JM, Loudy DE, Huelsken J, Birchmeier W, et al. beta-Catenin is required for specification of proximal/distal cell fate during lung morphogenesis. J Biol Chem. 2003;278(41):40231–8.

55. Chu EY, Hens J, Andl T, Kairo A, Yamaguchi TP, Brisken C, et al. Canonical WNT signaling promotes mammary placode development and is essential for initiation of mammary gland morphogenesis. Development. 2004;131(19):4819–29.

