## Supplementary material for "Inference and validation of an integrated regulatory network of autism": Supplementary Information 11.docx

| Phenotype Term | Involved tissues |
| --- | --- |
| Abnormal nervous system physiology | Central nervous system  Cranial nerve  Ganglion  Glymphatic system  Nerve  Nerve plexus  Neural crest  Neuroendocrine gland  Peripheral nervous system  Tail nervous system |
| Abnormal CNS synaptic transmission | Brain  Central nervous system ganglion  Central nervous system nerve  Central nervous system roof plate  Central nervous system vascular element  Ependyma  Floor plate  Future spinal cord  Gray matter  Meninges  Neural plate  Neuraxis flexure  Primitive meninges  Spinal cord  Tail central nervous system  Ventricular layer  White matter |
| Nervous system phenotype | Central nervous system  Cranial nerve  Ganglion  Glymphatic system  Nerve  Nerve plexus  Neural crest  Neuroendocrine gland  Peripheral nervous system  Tail nervous system |
| Abnormal brain morphology | Brain blood vessel  Brain ependyma  Brain floor plate  Brain gray matter  Brain mantle layer  Brain marginal layer  Brain meninges  Brain roof plate  Brain subventricular zone  Brain vascular element  Brain venous system  Brain ventricle  Brain ventricle and choroid plexus  Brain ventricular layer  Brain white matter  Brainstem  Choroid plexus  Circumventricular organ  Forebrain  Forebrain-midbrain boundary region  Hindbrain  Midbrain  Midbrain-hindbrain junction |
