## Supplementary figures and images for "Inference and validation of an integrated regulatory network of autism"

### Supplementary Information 1.pdf

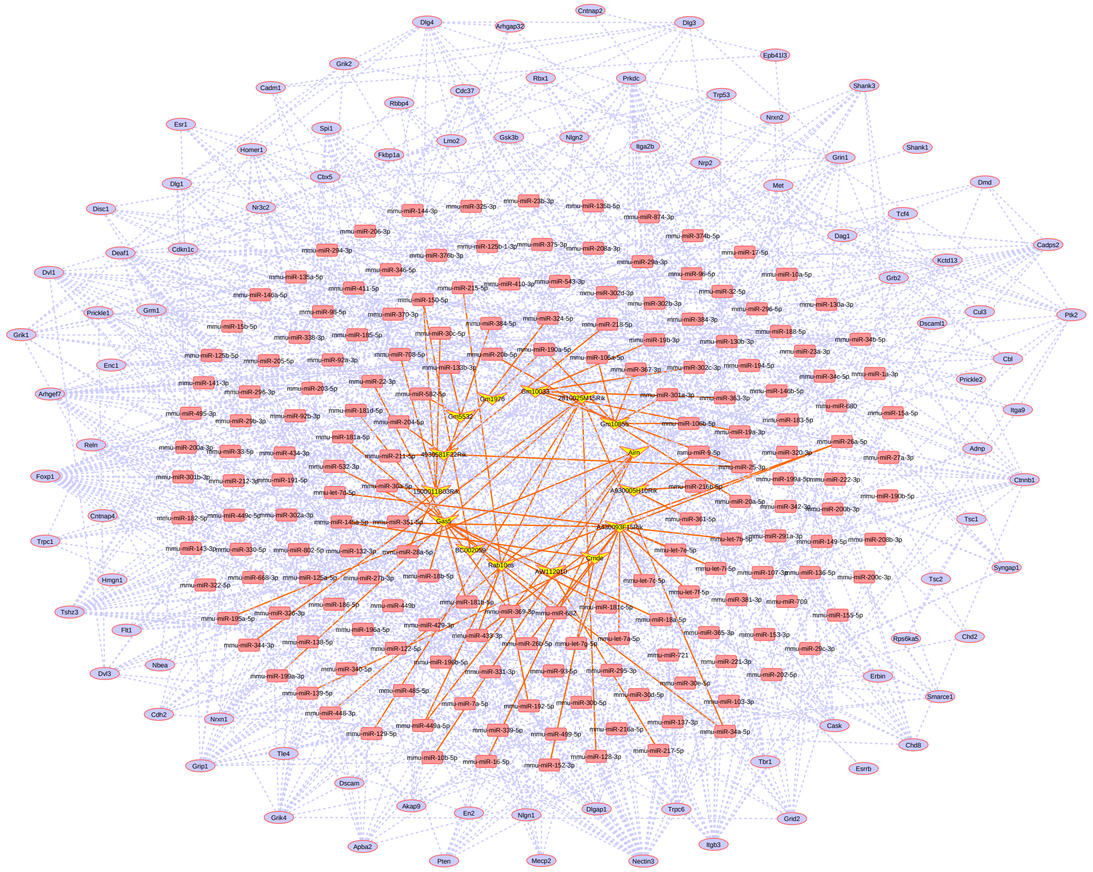

### Supplementary Information 4.jpg

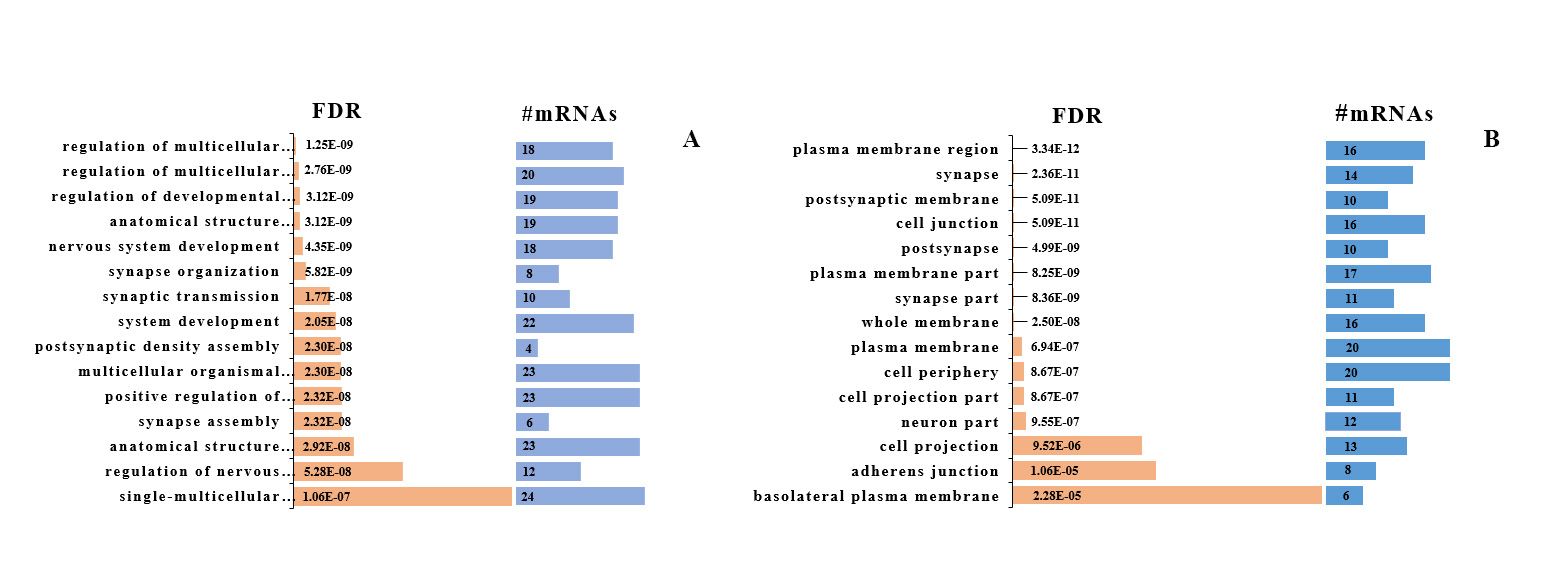

### Supplementary Information 6.jpg

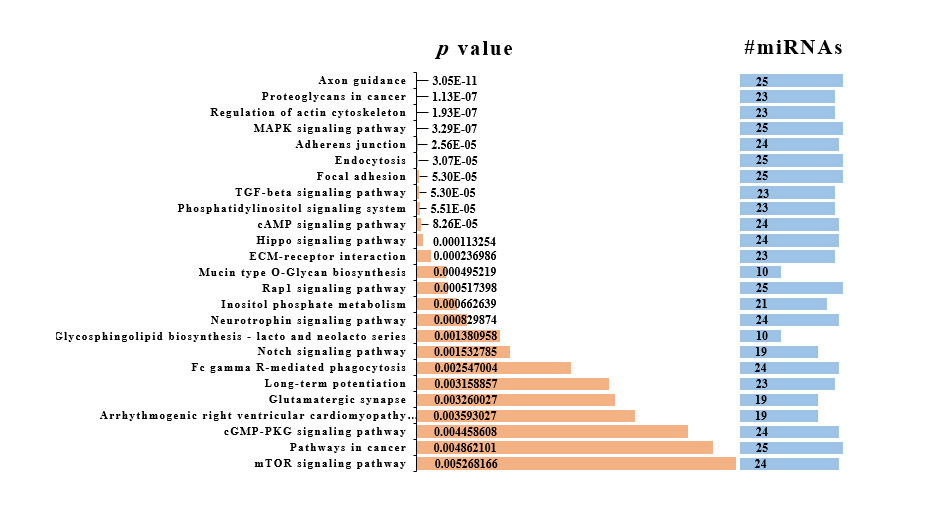

### Supplementary Information 7.jpg

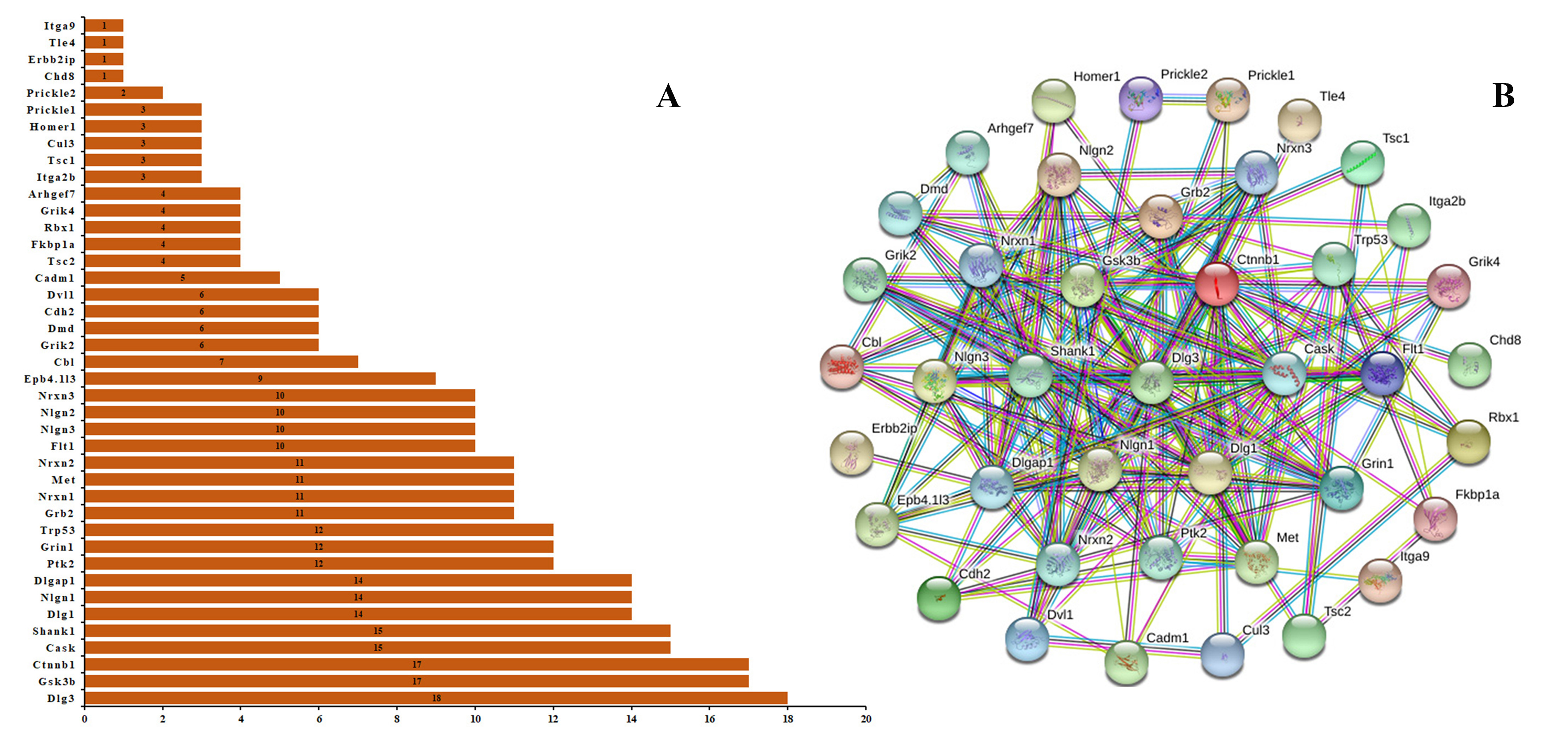

### Supplementary Information 10.jpg

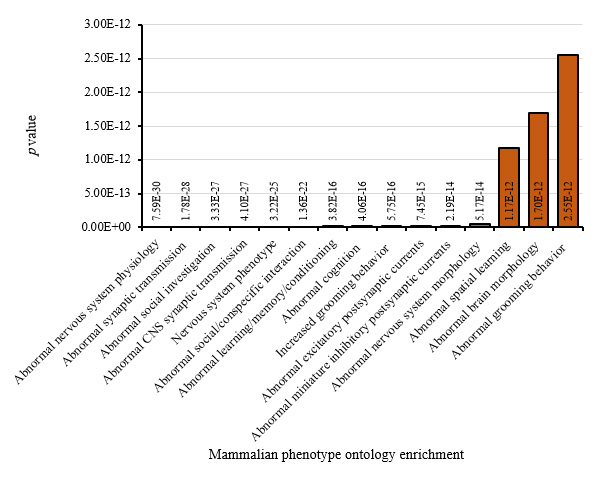

### Supplementary Information 12.jpg

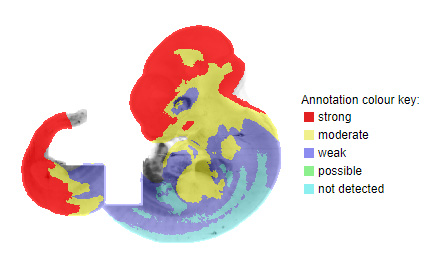

### Supplementary Information 13.jpg

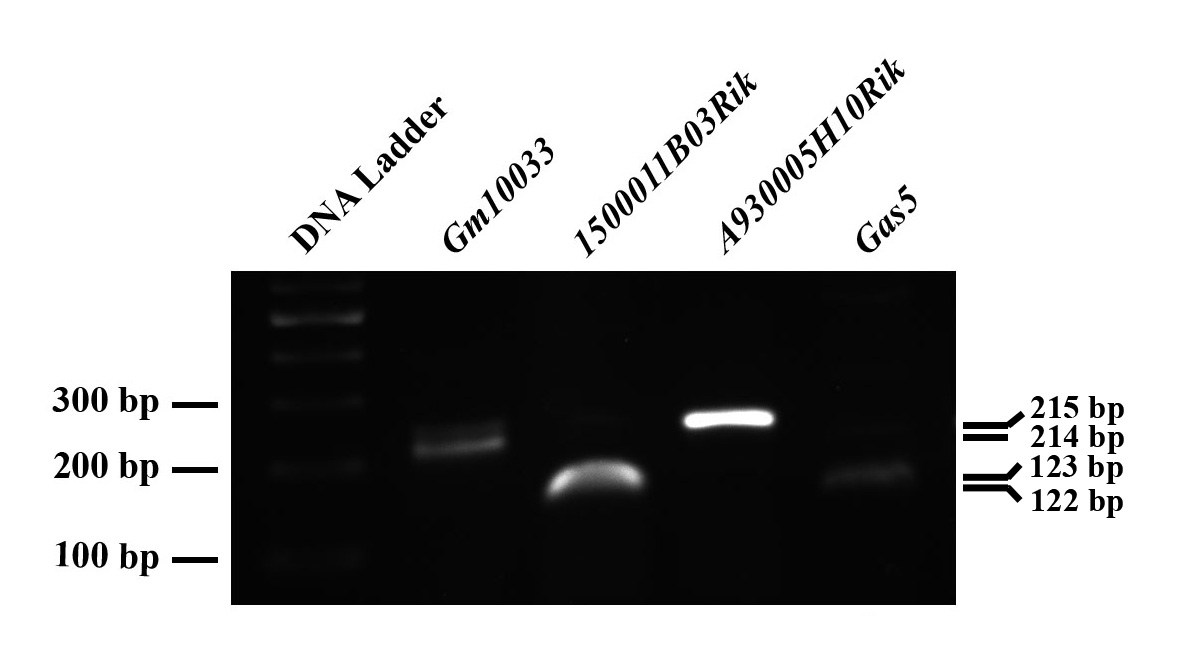

### Supplementary Information 14.jpg

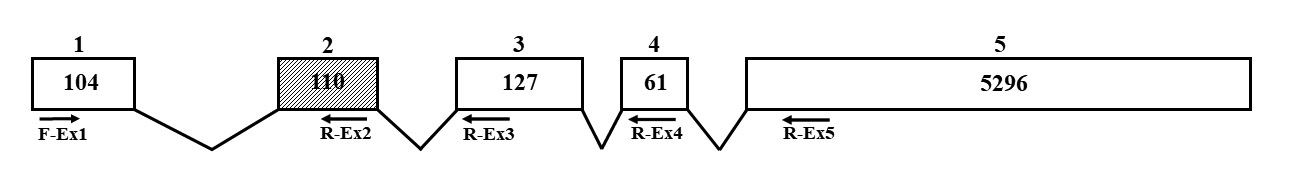

### Supplementary Information 15.jpg

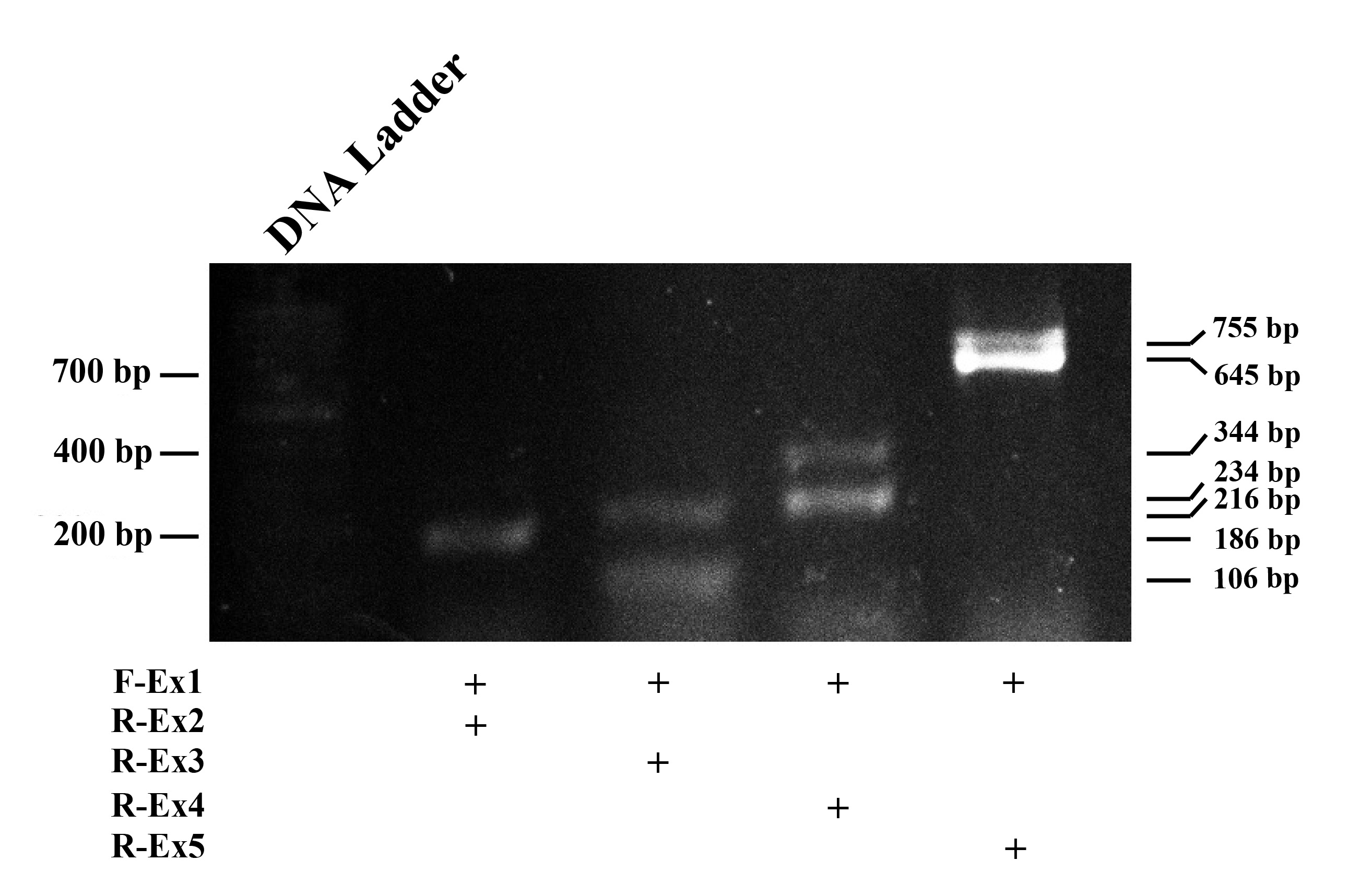

### Supplementary Information 16.jpg

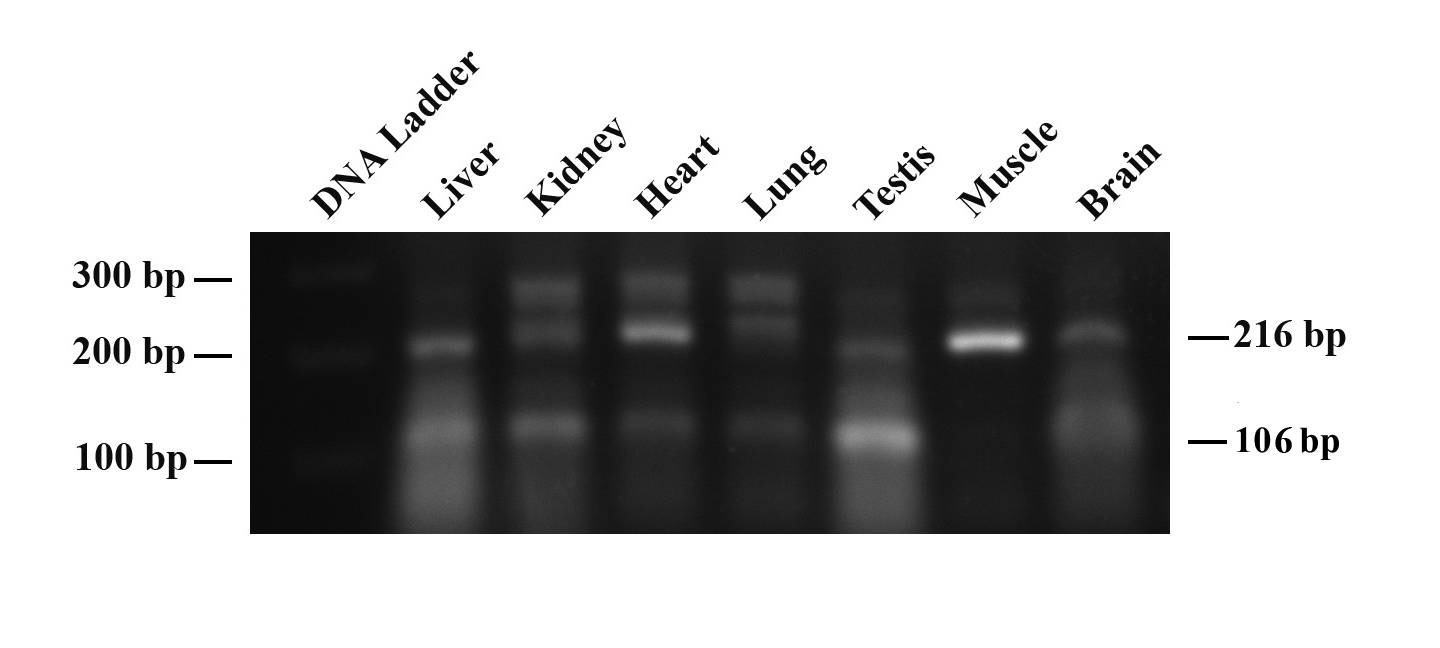

### Supplementary Information 17.jpg

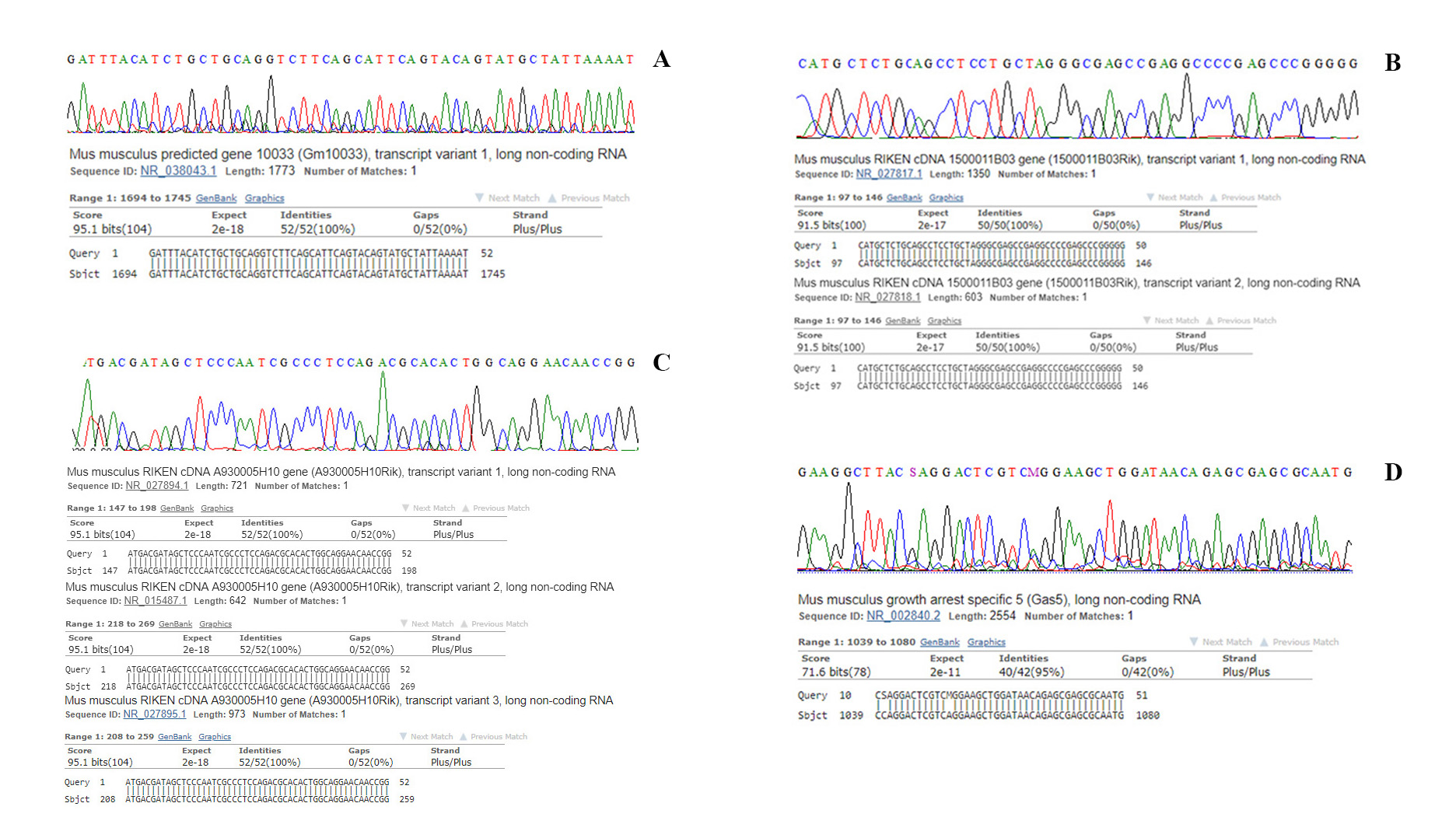
